# Detecting cultural evolution in a songbird species using community-science data and computational modeling

**DOI:** 10.1101/2023.01.23.525255

**Authors:** Yakov Pichkar, Abigail M. Searfoss, Nicole Creanza

## Abstract

Song in oscine birds is learned across generations, and aspects of the song-learning process parallel genetic transmission: variation can be introduced into both cultural and genetic traits via copy-error, and both types of traits are subject to drift and selective pressure. Similarly to allele frequencies in population genetics, observing frequencies of birdsong features can improve our understanding of cultural transmission and evolution. Uniquely, community-science databases of birdsong provide rich spatiotemporal data with untapped potential to evaluate cultural evolution in songbirds. Here we use both community-science and field-study recordings of chipping sparrows to examine trends across nearly seven decades of song. We find that some syllable types tend to persist in the population for much longer than others. Persistent songs tend to contain more syllables of shorter duration than songs that were observed across fewer years. To draw inferences about the effects of learning biases on chipping sparrow syllables, we construct a spatially explicit agent-based model of song learning. By comparing our empirical analysis to simulated song distributions using three different song-learning strategies—neutral transmission, conformity bias, and directional selection—we suggest that chipping sparrows are unlikely to select tutors neutrally or with a conformity bias and that they learn their songs with a remarkably low copy-error rate.

## Introduction

For oscine songbirds, song has many important functions, including territory defense, species identification, and mate attraction (Catchpole & Slater, 2003; Searcy & Andersson, 1986). In contrast to a closely related outgroup (suboscines), oscines must learn their songs, making the process of song learning critical to the reproductive success of individuals across this diverse clade (Kroodsma et al., 1982; Kroodsma & Miller, 1996; Mason et al., 2017; Thorpe, 1958). The evolutionary dynamics of learned song exhibit parallels to those of human cultural evolution, where long-lasting traditions can coexist (Aplin, 2016; Hoppitt & Laland, 2013; Kandler & Laland, 2009; Tomasello et al., 1993; Whiten, 2017). By studying song learning, we can better understand which aspects of human learning and cultural evolution are shared with other species and which properties are unique.

The transmission of information between individuals underpins both human and avian cultural evolution. Laboratory and field studies have shed light on how song is transmitted in avian populations. Some of these studies have measured the properties of cultural transmission: the similarity between learned song and tutor song, error rates in song matching, the invention of new songs, and the frequency of songs in a population, among other factors (Cardoso & Atwell, 2016; Marler & Peters, 1982; Marler & Tamura, 1962; Slater, 1986; Thorpe, 1958). Others have used field-site data to address questions of song change over time. For example, some studies have tracked cultural evolution using recordings taken in one population over multiple decades (Ju et al., 2019; Williams et al., 2013), and other studies have demonstrated that temporal changes in song are discerned by the current population by showing that birds react more strongly to modern recordings than to historical ones (Derryberry, 2007, 2011). Field-study recordings can ensure coverage of local song repertoires, facilitate direct observation of song tutors, and provide samples from the entire site’s population. Due to the limits on the time period and geographic range they can cover, field studies are snapshots of the cultural evolution of syllables, and larger-scale studies can help bridge the gap between local behaviors and cultural evolution..

By tracking songs and reproductive success over time, as well as by determining which song features correspond to stronger responses in current populations, researchers have gained insight into the types of selective pressures that operate on song (Derryberry, 2007, 2011; Williams et al., 2013). In parallel, evolutionary biologists and population geneticists, without access to time-series data, have synthesized evolutionary models with evidence from existing distributions of allele frequencies to understand whether regions of the genome have undergone selection (Bamshad & Wooding, 2003; Bustamante et al., 2001; Ford, 2002; Gutenkunst et al., 2009; Nielsen, 2005; Williamson et al., 2005). A genetic variant can become more frequent in a population because it is associated with a fitness advantage (selection) or due to random chance (genetic drift). A genetic region under selection will tend to have a different distribution of allele frequencies than those regions not under selection (Nielsen, 2005). Thus, one approach in population genetics is to simulate the evolution of a trait under different selection pressures and population histories. By comparing data from real populations to predictions from evolutionary models, researchers have identified which of these models best explains the data (Akashi & Schaeffer, 1997; Gutenkunst et al., 2009; Kryazhimskiy & Plotkin, 2008; Williamson et al., 2005). Some researchers apply this theoretical approach to the cultural evolution of song by examining the distribution of song within populations (Lynch et al., 1989; Lynch & Baker, 1993, 1994; Mcgregor & Krebs, 1982; Parker et al., 2012) and by developing individual-based or agent-based simulations of song learning that are compared to field-site data (Crozier, 2010; Ellers & Slabbekoorn, 2003; Lachlan et al., 2018; Lachlan & Slater, 2003; Slater, 1986; Wheelwright et al., 2008; Youngblood & Lahti, 2022). Such agent-based simulations have been used in conjunction with birdsong data to infer the learning strategies used by swamp sparrows and house finches (Lachlan et al., 2018; Youngblood & Lahti, 2022). With these comparisons of field recordings and results, researchers found evidence for different cultural transmission biases in different species. For example, swamp sparrows showed evidence of conformity bias, a type of frequency bias in which common song variants are disproportionately preferred (Lachlan et al., 2018). However, house finches showed evidence of content bias, in which certain song elements are preferentially learned regardless of their frequency in the population, a form of directional selection on the basis of a feature of the song (Youngblood & Lahti, 2022).

Here, we present an extension to this approach by developing a model of cultural transmission of birdsong and comparing the results of this model to a large-scale song analysis of community-science recordings. We suggest that utilizing community-science data is a time- and cost-effective supplement to field studies in the study of birdsong evolution. Specifically, community-science data—which can cover a large geographic area over many years—can provide a unique insight into patterns of song transmission across large spans of time or space, particularly when these data are considered alongside evolutionary models. Researchers have analyzed community-science data to examine avian behaviors, whereas other studies have compared spatially explicit models to song recordings from the field to examine evolutionary hypotheses (Bolus, 2014; Dennhardt et al., 2015; Goodenough et al., 2017; Kaluthota et al., 2016; Newson et al., 2016; Robinson et al., 2018; Silvertown et al., 2011). We synthesize these approaches by examining community-science data via models of song learning, providing insights into cultural evolutionary patterns.

As a focal species for this study, we chose the chipping sparrow, which has a simple repertoire of one repeated syllable. As a result, the full vocal repertoire of an adult bird can be captured by a single community-science recording. Since cultural transmission includes mechanisms of mutation, selection, and drift similar to those found in genetics, we employ techniques from population genetics—in particular, an adaptation of site frequency spectra (Bustamante et al., 2001; Nielsen, 2005)—to study song evolution. We identify unique syllable types that characterize the songs in a population, and use the occurrence and lifespans of these syllables to gain a deeper understanding of chipping sparrow learning. Since different learning strategies result in different distributions of syllables—many replicates of the same syllable persisting over time if birds learn based on a conformity bias, and potentially a relatively small number of syllables with desirable characteristics in the case of a directional bias—the frequency at which syllables occur in nature carries information about these biases. We compare the occurrence and longevity of songs to distributions produced by a computational model. This model simulates the transmission of syllables in a population under three types of learning— neutral evolution, conformity bias, and directional selection. We demonstrate how analysis of community-science data in association with a model can supplement field studies and extend the understanding of birdsong evolution.

## Methods

### Categorization of chipping sparrow syllables into types

In a previous study, we gathered and analyzed field-site and community-science recordings of chipping sparrows across the species’ entire breeding range (**Fig. S1**), measuring numerous acoustic features of each song and classifying the syllables into distinct types and categories (Searfoss, Liu, et al., 2020; Searfoss, Pino, et al., 2020). A number of recorded songs in our previous analysis (Searfoss, Liu, et al., 2020) did not have a recording date listed; however, by revisiting the original field recording notes we were able to find the years for all recordings for our study presented here. We categorized songs as follows: all songs were viewed as spectrograms in Audacity on a fixed frequency-and time-scale (see **Appendix S1**). A single syllable was then selected as representative of a song, since chipping sparrow songs are fully characterized by repetition of a single syllable. We manually classified 820 syllables into 112 chipping sparrow syllable types based on the shape of the syllable (**Appendix S1**, similar to methods of (Borror, 1959; Leitner & Catchpole, 2004; W.-C. Liu, 2001; Vargas-Castro et al., 2012); for examples of spatial syllable distributions, see **Fig. S15**). We further grouped these syllable types into broader categories based on the syllable shape: up-down (up-slur followed by down-slur), down-up (down-slur followed by up-slur), sweep (single up-slur or down-slur), complex (more than two slurs), doubles (a slur with multiple frequencies), and buzz (syllable containing a noisy and/or high-entropy section, generally termed ‘buzzy’). To ensure that we were correctly categorizing the repeated element, particularly in the case of up-down versus down-up syllables, we examined the beginning and end of the song to determine which part of the syllable came first.

In addition, we used the song-analysis software Chipper to extract eight song features from each recording: mean intersyllable silence duration, mean syllable duration, mean syllable frequency range, mean syllable minimum and maximum frequency, duration of song bout, mean syllable stereotypy, and total number of syllables. Chipper allows the user to visualize each song bout, and it predicts where syllable boundaries are located using fluctuations in the amplitude of the signal (Searfoss, Pino, et al., 2020). The user can change the signal-to-noise threshold, apply lowpass and highpass filters to exclude high-frequency and low-frequency noise, respectively, and manually correct these syllable boundaries if necessary. Then, Chipper analyzes the signal within each identified syllable and outputs a matrix of features for each song (see (Searfoss, Pino, et al., 2020) for more details on how each song feature is extracted from the spectrogram).

For the catalog numbers, database, recordist, URL, and license for the 820 song files, see **Table S1**. For the metadata including recording latitudes and longitudes and the 8 song features (all log transformed except mean stereotypy of repeated syllables and the standard deviation of note frequency modulation), see **Table S2**.

### Calculating and analyzing the lifespan of chipping sparrow syllable types

The observed “lifespan” of a syllable type was defined as the period between the earliest and latest years in which a syllable type was recorded. To explore the properties of these lifespans, we plotted the distribution of syllable lifespans and the number of times in which these syllables were identified. We proceeded to compare syllable features between songs that contained short-lived (recorded lifespan=1 year) versus long-lived (recorded lifespan≥50 years) syllable types. We performed Wilcoxon rank-sum tests on short-versus long-lived syllables for the eight song features extracted from each recording. For stringency, we conducted a Bonferroni correction for multiple hypothesis testing by dividing the *P*-value threshold for significance (𝛼=0.05) by the number of tests. Overall we performed one test on 8 song features, so the threshold for significance was lowered to 𝛼_adjusted_=6.25×10^-3^. In addition to our previously observed geographic patterns in chipping sparrow songs (Searfoss, Liu, et al., 2020), we also conducted Wilcoxon rank-sum tests to determine whether short-or long-lived syllables are more frequent on an east-west axis.

### Model design

We developed an agent-based simulation to model song learning in the chipping sparrow population. The entirety of the model was implemented in Python 3.7 and uses the following primary packages: NumPy v1.16.3, Matplotlib v3.0.3, Pandas v0.24.2, and SciPy v1.2.1 (GitHub link available on request but removed for double-blind peer review). With this model, we simulate syllable transmission in a population under three learning regimes—neutral transmission, conformity bias, and directional selection. Under a neutral model of song learning, a juvenile randomly chooses a tutor’s song to imitate; with conformity bias, a juvenile is disproportionately likely to choose a tutor with the most common song; and directional selection operates to favor certain song properties such as rate of syllable production or higher frequency bandwidth (Podos, 1997; Podos & Nowicki, 2004), such that juveniles are more likely to learn songs that exemplify better performance. Here, directional selection is somewhat analogous to the content bias observed in house finches (Youngblood & Lahti, 2022) since juveniles are choosing to learn a syllable based on its properties and not its frequency of occurrence. Chipping sparrows only learn a single syllable, so in our model, directional selection operates on a continuous feature of a syllable—the rate of syllable production—instead of the selection of certain syllable types to compose a song.

As illustrated in **Fig. 1**, we initialized each model with a 500×500 matrix of syllable types that represented a population of birds (we performed additional analyses with matrix sizes of 400×400, 600×600, and 700×700). Each matrix location represents a single bird that sings a single syllable—a categorical value that was initially assigned randomly from a discrete uniform distribution {1:500}. For the directional selection model, an additional matrix of identical size is used, containing continuous values representing a syllable feature (the rates of syllable production) randomly sampled from a truncated normal distribution confined to the values observed in nature, i.e. a minimum of 5 and a maximum of 40 syllables per second, with mean 22.5 and variance 25 syllables per second (*X∼N(22.5, 25), 5<X<40*) (Searfoss, Liu, et al., 2020). In each timestep, roughly corresponding to a year, the following steps take place: a fraction of birds die, juvenile birds are tutored and fill the empty territories, and a portion of the population disperses (**Fig. 1**). Note that there is a substantial burn-in period (discussed under ‘Sampling the simulated bird population’), so initial distributions of syllables and song-rates have minimal impact on final, sampled values.

**Figure 1.**
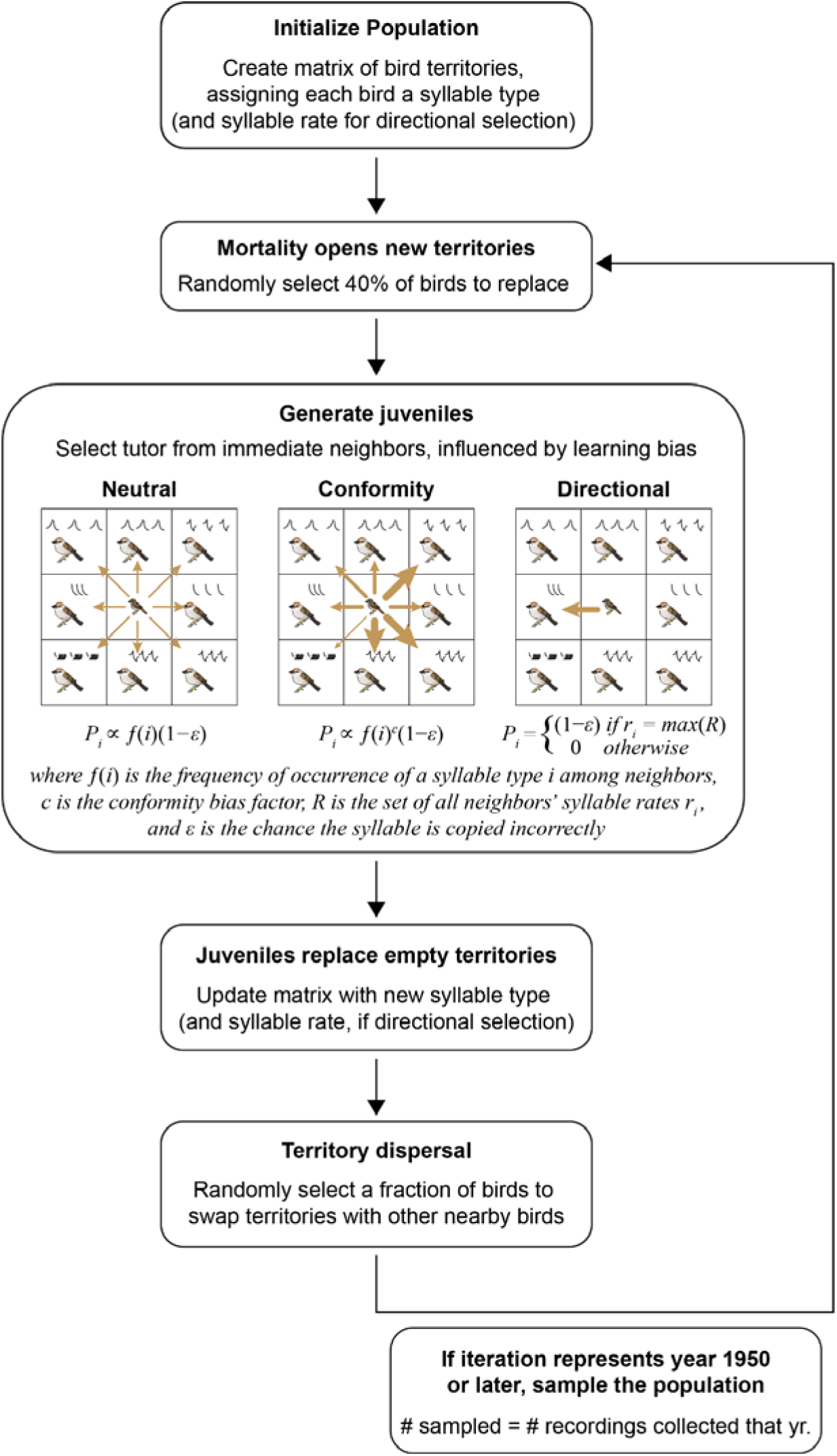
Model schematic with illustrated representation of learning biases. For neutral learning, the frequency of a syllable type among the juvenile’s adjacent neighbors is proportional to the probability that it will be learned by the juvenile (equal weight for all neighbors; identical arrows). Conformity bias modifies this probability by taking probabilities to the power of a conformity factor *c*, such that more common syllables are preferred. In our implementation of directional bias, syllable rate is the song characteristic that is selected for (although this could easily represent any song feature): the neighbor with the highest syllable rate is chosen as the tutor. Both the syllable type (for all learning models) and syllable rate are learned with some probability of error; if the syllable type is incorrectly copied, a new syllable is invented.

More specifically, in each timestep, first a fixed percentage of the birds are randomly selected for death. We set this mortality rate to 40% based on similar avian models (Lachlan et al., 2018; Slater, 1986). For every bird that is eliminated, a new juvenile bird replaces it; the new bird’s repertoire is either a novel syllable in the case of copy error, or a syllable learned from one of its neighbors (namely, the birds present at the beginning of the timestep and adjacent to its hatching location in the two-dimensional matrix—up to eight birds). In order to maintain a spatial arrangement representative of natural territories, the matrix boundaries do not wrap, so birds at the edges of the matrix have fewer neighbors. Maintaining father-juvenile relationships was not necessary in our model, as oblique transmission of song, rather than vertical song transmission from parent to offspring, appears to be predominant in chipping sparrows. In a well-studied population in the northeast US, juveniles learned a song before their first migration but often changed their song after migration to better match a neighbor, leading to a spatial pattern in which the syllable types of neighbors often differ but occasionally match closely; an individual’s song did not change further after the first year (W.-C. Liu & Kroodsma, 2006; W.-C. Liu & Nottebohm, 2007). (Although the phenomenon of post-migration song modification has not been studied in other chipping sparrow populations, the pattern of occasional neighbor-matching was also observed in chipping sparrows in Mexico (Marler & Isaac, 1960).) Once all juveniles learn a syllable (see next sections for the three learning strategies), the new syllable types replace the matrix elements of the birds that died, representing juveniles moving into vacant territories. Each new syllable was represented by a new integer, such that all syllable types could be uniquely identified. All territories vacated by birds that died are filled simultaneously, after tutor selection occurred for that time step. Since deaths in nature occur throughout the year and learning takes place over a short time, all birds present at the beginning of the timestep can influence the learning of juveniles during that timestep.

In addition, birds have an opportunity to disperse: some portion of birds (termed the ‘dispersal fraction’) is selected to move to a nearby location on the matrix. This promotes the mixing of regionally common syllable types with the larger population. Dispersal may sustain the local syllable diversity seen in chipping sparrow populations (W.-C. Liu & Kroodsma, 2006). The addition of this dispersal step reflects what has been observed in the field: adults occasionally move to a new location, especially when they share a song with a neighbor (W.-C. Liu & Kroodsma, 2006). We tested dispersal fractions between 0 and 1 in 0.1 increments, where 0.5 means that half of birds attempt to swap places with a bird that has not yet changed places, chosen randomly from a location within a set radius (up to 11 matrix units). These dispersal fractions and radius values were informed by field studies describing chipping sparrow territory size and dispersal patterns (W.-C. Liu, 2004; W.-C. Liu & Kroodsma, 2006; Swanson et al., 2004). For each set of parameters, the simulation is run for 1000 timesteps, of which the final 68 are used for sampling (to compare with the 68 years of available community-science data) and the first 932 are a long burn-in period prior to sampling.

### Model implementation of neutral tutor selection

During each timestep, birds that die are replaced by a juvenile bird at each location. For neutral tutor selection, the syllable type learned by this juvenile is chosen at random from its eight immediate neighbors (or fewer, at edges and corners). To account for some likelihood of the new bird producing a novel syllable, we include a probability of error in learning. The probability of learning syllable type *i* in the case of neutral tutor selection can be represented as

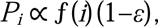

where ƒ(*i*) is the frequency of occurrence of a syllable type *i* among neighbors and ε is the chance the syllable is copied incorrectly. We varied this error rate parameter (𝜀) to explore the ranges of plausible error rates for each learning model (10^-6^ %, 10^-5^ %, 0.0001%, 0.001%, 0.01%, 0.1%, and 1.0%). We also ran several models in which an error meant that, instead of inventing a novel syllable, a bird produced a random syllable from the original set of syllables {1, 500}. For these models of syllable reinvention, called homoplasy, we added a larger error rate of 10%. Since only about 0.5% of observed songs were recorded only once (see **Fig. 2**), we did not test values of new syllable invention higher than 1%.

**Figure 2.**
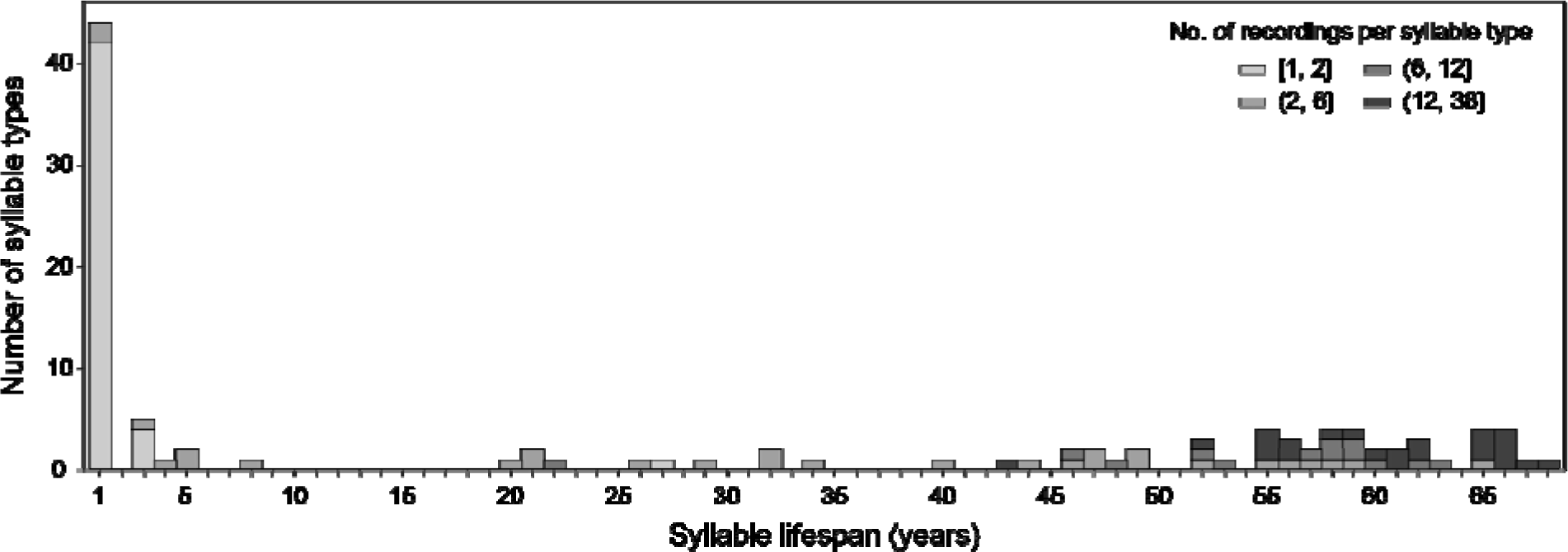
Distribution of chipping sparrow syllable type according to their lifespans. For our database of 820 recordings of 112 syllable types, we plot the number of syllable types versus syllable lifespan across the entire range. Each syllable type is also shaded by the total number of recordings of that syllable type, illustrating that longer-lived syllable types are also more common, though less common long-lived syllables exist.

### Model implementation of conformity bias

Under conformity bias, juveniles preferentially learn more frequent songs. Each juvenile surveys the syllable types sung by his neighbors. The probability of learning syllable type *i* in the case of conformist tutor selection can be represented as

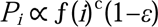

where *c* is the conformity bias factor. In the simulations we report below, a conformity factor *c*=2 means that each syllable’s frequency is squared. These values are then normalized to represent the likelihood of selecting each syllable, such that more common syllable types are learned more often than they appear among neighbors. We tested a series of other conformity factors including less severe conformity biases (*c*={1.2, 1.4, 1.6, 1.8, 2.0}) and a weak novelty bias (*c*=0.8). The learning error and dispersal were examined identically to those of the neutral tutor selection model.

### Model implementation of directional tutor selection

For directional tutor selection, learning is based not on the frequencies of syllable types, but on a continuous variable representing the rate of syllable production. The probability of learning syllable type *i* under directional selection is given by

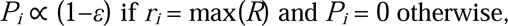

where *R* is the set of all neighbors’ syllable rates *r_i_*. This process mimics a type of directional selection that has been proposed for chipping sparrows, in which the preferred song is most difficult to produce. This model could accommodate directional selection on any continuous song feature; as a case study, we use a simple metric—rate of syllable production, also called trill rate—as our putative feature that is under selection. This feature has been hypothesized to be relevant in chipping sparrows (Goodwin & Podos, 2014), but we note that our previous analysis showed a relatively wide distribution of trill rates in the chipping sparrow population over time, from ∼5 to 40 syllables per second. In the model, the juvenile selects a tutor—the neighbor with the fastest syllable rate—and attempts to copy both the syllable type and the syllable rate of this tutor. The learning error for syllable type operates identically to the neutral and conformity models. With directional selection, there is also a learning error for syllable rate which is weighted such that it is more difficult to replicate or improve upon the tutor’s performance than to perform worse than the tutor (as in (Henrich, 2004)). Thus, juveniles are most likely to sing at a slightly slower rate than the tutor, and the syllable rates are restricted to values observed in our chipping sparrow song database (between 1 and 40 syllables per second). Therefore, the learned song is sung at a rate similar to that of the tutor with an error drawn from a uniform distribution between −2 and 0.25, and has either an identical or a novel syllable type. It is important to note that while we describe the song feature under selection as ‘syllable rate’, it can be interpreted that any continuously varying song feature is undergoing directional selection.

### Sampling the simulated bird population

The method of data collection from the model population was chosen to replicate the sampling that occurred when songs were recorded by community-scientists. For each learning model, the simulated population is sampled such that the number of birds sampled per timestep is equal to the number of recordings we have from each year. Our recording data spans 68 years (1950–2017) with some years having no recordings; thus some number of birds (possibly 0) are selected at random from the model population for each of the last 68 of 1000 iterations of the model (here, the first 932 timesteps serve as a burn-in, which minimizes the effects of initialization for variables such as the number and distribution of syllables). For each bird sampled from the simulated population, the iteration from which it was collected and the syllable type were documented. This was to ensure the lifespans and the counts of syllable types of this sampled model population could be calculated in the same manner as the recording data.

We sampled syllables from our simulations to compare the model to three similarly sized regions of the range chosen for their high sampling density. For regions of approximately 100,000 km^2^, we chose three regions with the highest density of song recordings. These regions were defined as the area within a rectangle bounded by latitudes and longitudes; these included the Michigan/Ohio region (85**°**W to 82**°**W and 39**°**N to 43**°**N, containing 172 song recordings over ∼100,000 km^2^), the New York region (77**°**W to 73**°**W and 40**°**N to 43**°**N, containing 88 songs over ∼100,000 km^2^), and the New England region (74**°**W to 69**°**W and 41**°**N to 45**°**N, containing 210 songs over ∼130,000 km^2^). We estimate that the chipping sparrow population density in the US to be about 13.3 birds per km^2^, calculated as the population size of chipping sparrows (estimated as 100 million chipping sparrows in the US (Will et al., 2020)) divided by the area of the continental US (∼7.5 million km^2^). We assume that our sampled regions likely have at least this average density, so we used a range of 13.3 to 25 birds per km^2^ for our calculations. For a breeding region of 100,000 km^2^, we thus estimate a population size of about 1.3 to 2.5 million chipping sparrows. Of these, approximately half are males, and not all males are adults with territories: likely only one in three chipping sparrows is a territorial male. Given their sizes, each region likely has 444,000 to 833,000 territories, compared to the 160,000 territories in the smallest simulations (400×400) and 490,000 in the largest (700×700). All these regions are concentrated towards the east of the chipping sparrow range, as equivalently sized regions in the west had a much smaller number of recordings (22 in California’s Central Valley, and fewer in Oregon). To estimate the syllable lifespans and counts that each model produced for each region, we sampled syllable types from every model 50 times.

### Quantitatively comparing model results to empirical data

For each simulation, we visualized and compared our results using the site-frequency spectrum technique from population genetics (Nielsen, 2005; Pepperell et al., 2013; Zhu & Bustamante, 2005) and created a “syllable frequency spectrum”—the frequency of birds that sing various syllable types in the sample—to compare our model results with empirical data. Additionally, we use a similar visualization to compare the frequencies of syllable-type lifetimes for each model across a range of learning error rates.

Specifically, we aimed to identify whether one or more tutor-selection models would be able to produce results similar to the empirical data, across both syllable frequency of occurrence and syllable lifetime distributions. These distributions have similarities to the empirical data: there are several frequent long-lasting syllables, many syllables observed very few times, and a small number of syllables with intermediate observed lifespans (**Fig. 2** for lifespans from the entire range). These intermediate-lifespan syllables were most difficult to replicate among these song-learning models in a way that could be detected by direct comparison of distributions, such as via two-sample Kolmogorov-Smirnov and *k*-sample Anderson-Darling tests; the sparsely populated intermediate values of syllable frequency and lifespan contained most of the relevant differences between models, whereas most mass used in comparisons of distributions lay at the extremes (**Figs. 2 & 5**). Instead of manually assigning categories of *short-longevity*, *intermediate-longevity*, etc., we placed the empirical data into bins using an algorithm; it assigned bin edges by minimizing the combined variance of bin count and bin width. First, we placed syllables that were recorded only once (for the syllable frequency spectrum) or only in a single year (for the distribution of syllable lifespans) into their own category, and all others were initially placed into six equally spaced bins. Then, the edges of these bins were progressively shifted 10^6^ times (the edges were moved, and variances of the resulting bins’ sizes and their number of data points were calculated). If the combined variance was lower, this became the new set of bins for the next permutation (see GitHub for details).

In this way, we used an algorithm to place the empirical chipping sparrow data into bins by syllable type using the optimally found edges, and the simulated samples were placed into the same bins to compare their distributions with the empirical data. On the binned syllable-type and lifespan spectra, we conducted Fisher’s exact tests between the empirical data (null hypothesis) and all simulated datasets to determine which combination of learning strategy, learning error, and dispersal fraction results in syllable-type and lifespan distributions that are concordant with the patterns found in the chipping sparrow population.

### Ethical Note

All recordings were gathered from open-access databases of recordings or published studies from the wild. Playback may have been used to elicit these songs, which can affect bird behaviors in the wild (Harris & Haskell, 2013). The short- and long-term effects of field playbacks are understudied and likely differ by species, but the limited use of playbacks (i.e. when precautions are taken not to unduly disturb birds and relatively few playbacks are presented at a reasonable volume and duration) is thought to be an important part of an ethical birdwatching practice (Sibley, 2011; Watson et al., 2019). Recordists uploading to community-science repositories are encouraged to document their use of playbacks, and few of our recordings were noted to be collected in response to playback.

## Results

### Syllable types and cultural analysis

We categorized syllables from 820 recordings into 112 distinct syllable types (**Appendix S1**, also see previous analysis (Searfoss, Liu, et al., 2020)). We found that syllable types that continue to exist for much longer than the lifetime of a chipping sparrow (less than 9 years) are also those that are most commonly observed, whereas other syllables are transient, and are observed rarely (**Fig. 2**). In other words, we did not observe any syllables that were found in many recordings but existed for a short period of time. To identify differences between long- and short-lived syllables, we classify syllable types as short-lived (lifespan=1 year) or long-lived (lifespan≥50 years), and compare the features of these syllables (**Fig. 3**). We find that long-lived syllable types are significantly shorter in duration than short-lived syllables (*P*<3.29×10^-3^) (**Fig. 3A**, **Table 1**), and songs with a long-lived syllable type contain significantly more repetitions of that syllable per bout (*P*<1.33×10^-3^) (**Fig. 3B**). We also found that geographic differences did not explain the trends in longevity, since the longitude of recordings of different syllable types did not significantly differ between the two lifespan categories (*P*=0.844). Additionally, buzz syllables tend to be long-lived, whereas double or complex syllables tend to be short-lived, with up-down, down-up, and sweep syllables being prominent in both lifespan groups (**Fig. 3C**).

**Figure 3.**
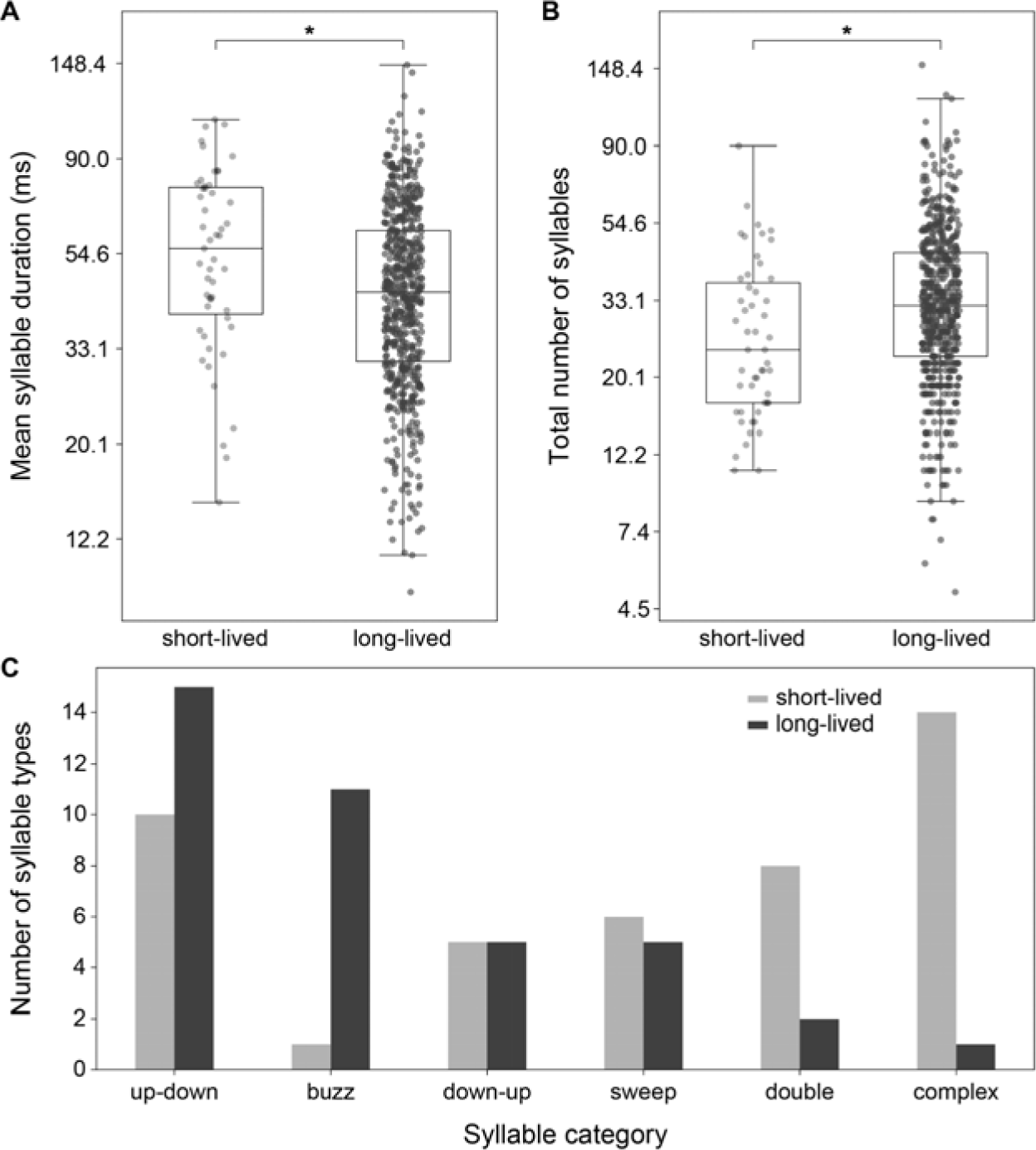
Syllable categories and song properties of short- and long-lived chipping sparrow syllabl types. Songs with long-lived syllable types (>50-year lifespan) have (A) significantly shorter syllables (*P*=3.29×10^-3^) and (B) significantly more syllables per song (*P*=1.33×10^-3^) than songs with short-lived syllable types (1-year lifespan). Significant results are indicated (* denotes *P<*6.25×10^-3^ from Wilcoxon rank-sum tests). (C) The number of short- and long-lived syllable types in each syllable category.

**Table 1.** Results of the Wilcoxon rank-sum tests between short- and long-lived syllable types.

| Song Features | Short- vs Long-Lived *P*-value |
| --- | --- |
| Mean intersyllable silence duration | 0.449 |
| Mean syllable duration | **0.003** |
| Mean syllable frequency range | 0.802 |
| Mean syllable minimum frequency | 0.476 |
| Mean syllable maximum frequency | 0.475 |
| Duration of song bout | 0.010 |
| Mean stereotypy of repeated syllables | 0.653 |
| Total number of syllables | **0.001** |

Bold indicates $P < 6.25 \times 10^{-3}$ .

### Model results

Here, we ran several models of song transmission in order to test which model produces patterns most similar to empirical data. Since the entire chipping sparrow range contains approximately 240 million individuals (Will et al., 2020), we compared the distributions of syllable occurrences and syllable lifetimes from our model to several focal regions that had high rates of sampling. We conducted a series of parameter sweeps, running the simulation for each of the three learning strategies with multiple error rates (or invention rates) spanning 10^-6^% to 1.0% and with territory dispersal fractions spanning 0 to 1 (i.e. 0 to 100% of individuals swap territories every timestep with a bird that has not already dispersed). We sampled the simulated syllables at the same frequencies as in the empirical data. We quantitatively determined which combinations of parameters (song-learning strategy, learning error rate, and dispersal fraction) produced results that did not deviate from the null hypothesis (empirical data).

We provide the syllable type and syllable lifetime frequency spectra for the best-fit model for each learning type. We determined a ‘best-fit’ model by first ranking the *P*-values (from least to greatest) for both the analysis of syllable type frequencies and the analysis of syllable lifespans and then combining the two ranks for each parameter set and choosing the one with the highest aggregate ranking, i.e. the largest sum of the two rankings (**Table S3**, **Fig. 4**). (We note that the results of the Fisher’s exact test indicate that the model results are potentially drawn from the same distribution as the real data when the null hypothesis is *not* rejected; when the *P*-value is not significant, the results of the model are statistically indistinguishable from the empirical data.) For one of our focal regions, New York, we find that the best-fit model for each learning type was: neutral tutor selection with 10^-5^ % learning error and 0.3 dispersal fraction (syllable frequency distribution *P*=5.7×10^-13^, lifespan frequency distribution *P*=9.5×10^-6^), conformity bias with 10^-3^ % learning error and 0.1 dispersal fraction (*P*=4.5×10^-8^ and *P*=8.2×10^-4^ respectively, with conformity factor *c*=1.2), and directional selection with 10^-6^% learning error and 0.5 dispersal fraction (*P*=0.0044 for occurrence spectra and statistically indistinguishable from the empirical lifetime data, *P*=0.123) (**Fig. 5**, see **Fig. S2** for corresponding unbinned spectra). For the other two focal regions, the best-fit parameters for each model were similar (**Table S4**, for spectra see **Fig. S4**, for parameter sweeps in other regions see **Figs. S9-S11**).

**Figure 4.**
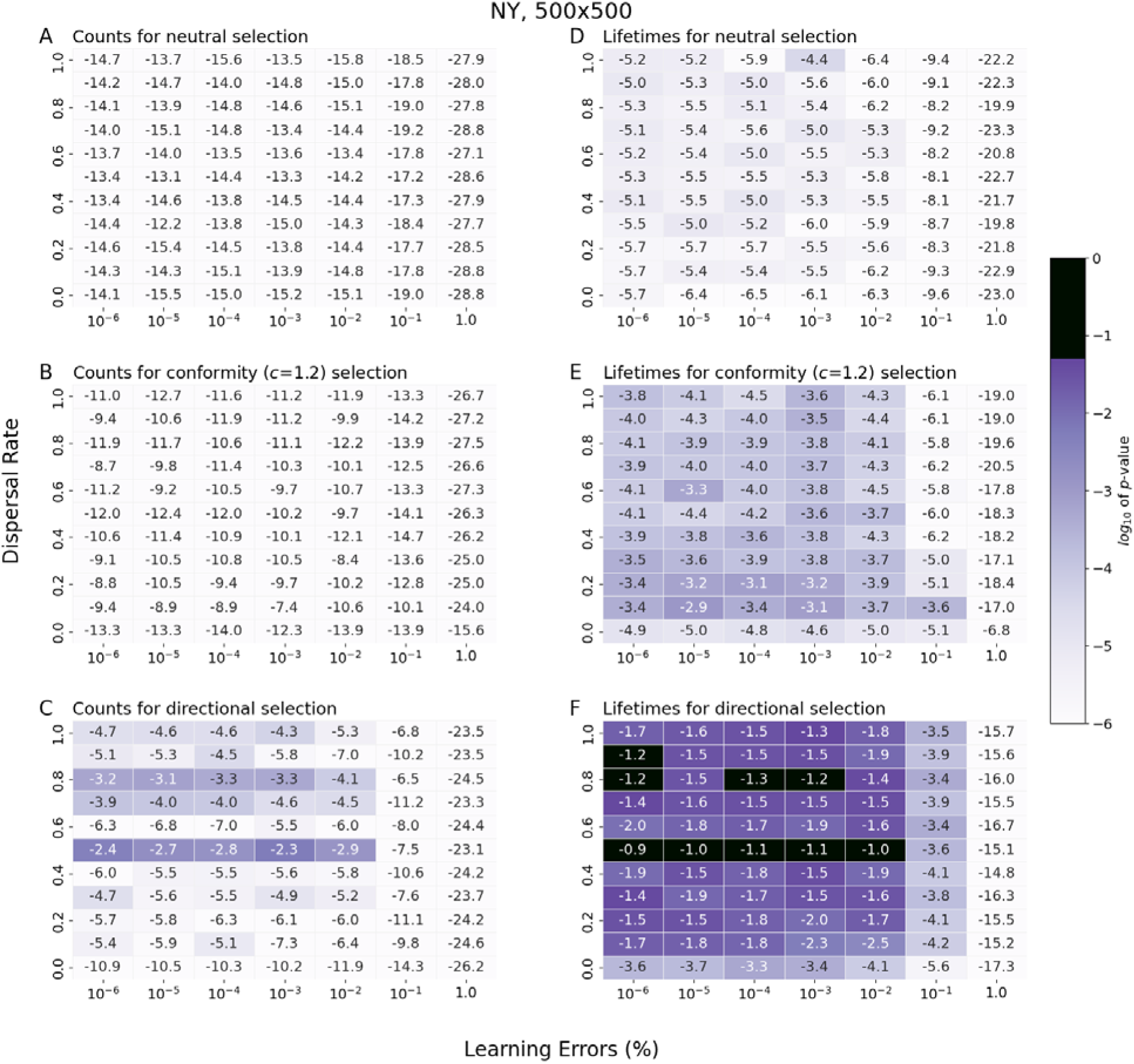
Statistical comparison of syllable frequency (A-C) and lifetime spectra (D-F) between computational models and empirical data. Sets of parameters (learning error and dispersal rate) for which the models and empirical data produce similar spectra distributions have *P*-values greater than 0.05 (shown in black). All simulations with neutral learning (A & D), conformity bias (with conformity factor *c*=1.2) (B & E) produced results that were statistically different from the empirical data (*P*<0.05). Some simulations with directional selection (C & F) produced lifetime spectra statistically indistinguishable from the empirical data (F) (*P*>0.05), though this was not true for occurrence spectra (C). For other regions, see **Fig. S6**.

**Figure 5.**
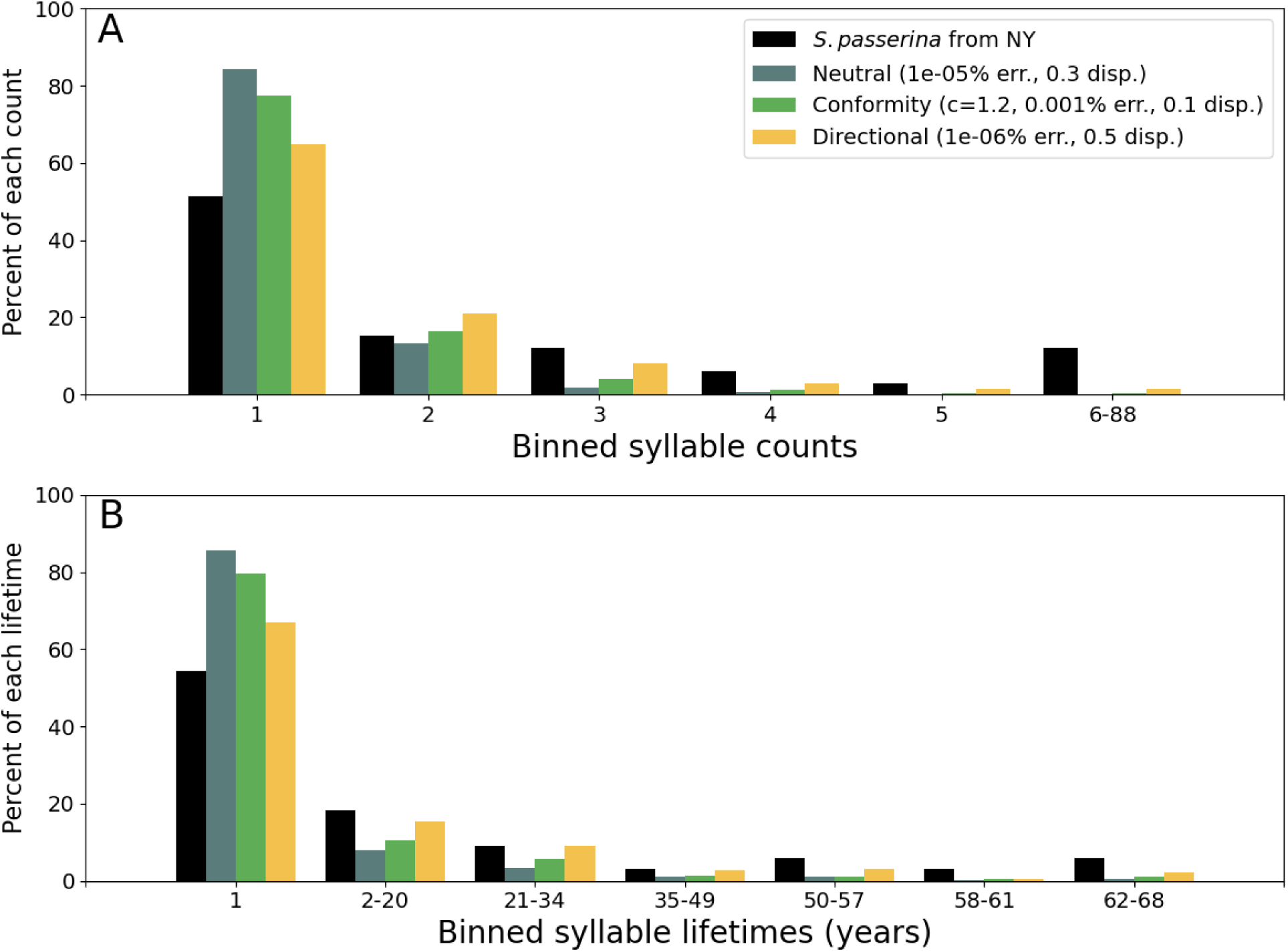
Comparison of binned syllable frequency and syllable lifespan spectra between empirical data and samples from best-fit models from one focal region. The number of times a syllable was sampled (A) and syllable lifetimes (i.e. 1 + (the last year) − (the first year in which the syllable type wa sampled)) (B). Each panel includes data from the best-fit models of each of the three song learning strategies: neutral tutor selection (blue), conformity bias (green), and directional selection (yellow). Data from community-science data is also provided (black). (See Methods for how bins were calculated.) For unbinned data, see **Figs. S2-3**; for frequency and lifespan spectra from other regions, see **Figs. S4-5**.

We measured the effect of changing several model parameters, including the strength of conformity selection, matrix size, and syllable reinvention. To measure the strength of conformity bias during selection, we varied the conformity factor *c*. Neutral selection (equivalent to *c*=1) and strong conformity selection (*c*=2) performed poorly, whereas an intermediate strength of conformity selection (*c*=1.2) performed best (**Figs. S7-8**, **Table S3**). However, even the simulation of conformity biased learning with the best performance (*c*=1.2, 10^-3^ % learning error and 0.1 dispersal fraction) produced distributions of syllable occurrences and lifetimes dissimilar to those observed in any region. To observe how varying the modeled population would affect the distributions of syllables, we simulated several population matrix sizes. We find that the larger the population matrix, the more difficult it is for our model to approximate the syllable lifetime distributions derived from the community-science data (**Table S5,** panels G & H in **Figs. S13-S15**), and that smaller matrices reproduce the empirical syllable lifetime distribution at a wide array of values (panel E in **Figs. S13-15**). Finally, we modeled whether homoplasy, the reinvention of syllable types, affects these distributions. Since homoplasy could only occur as the result of an error in song learning, the differences between models with and without syllable reinvention were greatest when error rates were high (**Figs. S9-12)**. The best-fit models with homoplasy had higher dispersal rates (**Figs. S9-12, Table S6**), but models with homoplasy did not describe the observed parameters better than models without homoplasy: models with high learning error—those that homoplasy affected the most—did not fit the empirical data for any combination of parameters.

Our regional models could not reproduce both the syllable counts and lifetimes found in the empirical data (**Fig. 4A-C**, **Table S3-4**). The best-fit model (of those with a matrix size of 500×500 territories), which relied on directional selection, low learning error (10^-6^% in NY and OH/MI, and 10^-4^% in New England) and intermediate dispersal fraction (0.5 in all regions), only reproduced the lifespans found from the community-science sampling (*P*=0.123 in NY, *P*=0.076 in OH/MI, and *P*=0.055 in New England, **Fig. S4** and **Table S4**). These models did not reproduce the empirical occurrence spectrum in any region (*P*=0.0044 for NY, 0.0037 for OH/MI, and 1.4×10^-4^ for New England) (**Fig. 4**, **Table S4**). The simulation with directional selection of tutors appears to most closely match the frequency of syllable types found in our empirical data, similar to the distribution of singletons and a long, flat tail (**Fig. S3** panels E-H). While long-lived syllable types arise in all three song-learning models, directional selection has an enrichment for these syllable types compared to the neutral model, whereas conformity bias has an abundance of long-lived syllables and few syllables with intermediate lifespans; thus, directional selection’s frequency spectrum of syllable-type lifespans best reflects what is observed in our community-science data sample (**Fig. 5B**).

## Discussion

Here, we performed an analysis of chipping sparrow song recordings across nearly seven decades to identify long-term patterns. We extended the use of computational approaches to cultural evolution (Youngblood, 2019) with techniques from population genetics and rapidly growing community-science data (Searfoss, Liu, et al., 2020) to assess cultural change and stability in birdsong. Community-science recordings provide broad spatiotemporal coverage of a species’ range, resulting in a dataset in which temporal changes could be identified across the entire population. By comparing these rich datasets with the predictions of cultural evolutionary models, we were able to evaluate the possible strategies underlying the social transmission of song. Specifically, we constructed a spatially explicit model of cultural transmission of chipping sparrow songs with different types of learning bias—neutral evolution (unbiased transmission), directional selection (favoring a certain characteristic of song), and conformity bias (favoring locally common songs). By comparing our empirical and simulated results, we found that a directional model most closely replicates the patterns of both syllable longevity and counts produced by chipping sparrow song learning. In addition, our computational analyses agreed with evidence from field research in finding that chipping sparrows have high-fidelity song learning (predicted new syllable invention rate of less than 0.1% in focal region analyses) and likely disperse to new territories (a dispersal rate of 0.1 or greater, most likely near 0.5) after initial learning (**Table S4**, **Figs. 4F & S6**).

Past studies have examined the diversity in syllables within the chipping sparrow population. For example, in the 1950s, Borror classified chipping sparrow syllables into categories and further subdivided the 58 recordings into 28 syllable types, demonstrating great song diversity and few observations of each syllable type (Borror, 1959). In a later analysis, the syllables of 157 chipping sparrows from the Eastern United States were analyzed and placed into around 30 distinct syllable types by eye (W.-C. Liu, 2001). With 820 songs, we identified 112 syllable types (**Appendix S1**). While our larger community-scientist-informed sampling is far smaller than the current chipping sparrow population—which is on the scale of 240 million—our analysis contains syllables that exist for decades (in the entire range **Fig. 2** and in focal regions **Figs. 5B, S4E-F**) and captures variation in birdsong that could not be identified via field studies of a species with such a large range (**Fig. S16**). It is possible that the sampled chipping sparrow syllables appear identical by chance rather than common descent as a result of syllable reinvention—a sort of cultural homoplasy. However, when we modeled syllable reinvention, models with homoplasy do not fit the empirical data better than models without it for any form of selection (**Table S6, Figs. S9-12** comparing panels A-F to panels G-L). Given the high fidelity of pupil learning (W.-C. Liu & Kroodsma, 2006), the presence of geographically clustered syllables (**Fig. S16**) (Searfoss, Liu, et al., 2020), and the results of our models, learning errors rarely result in birds reproducing an existing syllable elsewhere in the chipping sparrow range. These data have allowed us to explore trends in chipping sparrow song over time that will inform future studies of their song and of cultural evolution.

With our analysis of chipping sparrow syllables sampled from their entire range, we found that many syllable types appear to be rare and short-lived, whereas others are quite common and can persist for decades (**Fig. 2**). This was true both for the entire region, and when dividing the entire range into focal regions (**Figs 5B, S4E-F**). Further, we found evidence that some broad characteristics were associated with longer syllable lifespans. Buzz syllables tended to be long-lived whereas complex syllables tended to be short-lived, and songs with long-lived syllable types had more repetitions of shorter syllables, which would be consistent with predictions that songs with faster syllable repetitions might be favored in birds (Byers et al., 2010). Notably, this pattern of shorter and faster syllables being long-lived is geographically distributed: the distribution of short-versus long-lived syllables was independent of longitude (Wilcoxon rank-sum test, *p*=0.992) despite songs having more, shorter syllables on average than songs in the Western U.S./Canada than in the Eastern U.S./Canada (Searfoss, Liu, et al., 2020). Our results demonstrate that the diversity of chipping sparrow syllable types was not fully sampled in previous studies, and it is likely that other syllable types will be discovered as contributions of song recordings to community-science databases become more widespread. These results raise an important question: are syllables common and long-lived because of neutral transmission (similar to genetic drift), culturally favorable properties (i.e. certain syllables are inherently salient or associated with successful birds), or conformity bias (i.e. common syllables are preferred when learning song)?

Selectively neutral processes of song learning, such as unbiased learning of a song with a relatively high rate of error, are predicted to result in a simple pattern of syllable prevalence: most sampled syllable types would be sung by only one bird, fewer syllables would be sung by two birds, even fewer by three birds, and so on, until only a small handful of syllables might be sung by many birds (Slater, 1986). Slater observed this distribution of syllables in chaffinches: in a population of 36 chaffinches, most songs were sung by only one bird, but one song was sung by 22 birds. Further, he modeled the song-learning process with a simulation in which newly settled birds learned a random nearby song with some error; this simulation demonstrated that a neutral learning process with a predictable rate of copy-error was sufficient to replicate the observed distribution of chaffinch syllables. A similar pattern is regularly observed in genetic data in a stable population in the absence of selection pressures: most genotypes are rare, and few genotypes predominate (Nielsen, 2005). Thus, for both genotypes and song types, one does not need to invoke selection pressures to explain a pattern in which one or very few types are widespread but most are observed only once.

The question of whether directional selection plays a strong role in chipping sparrow song evolution has been a topic of debate in the literature (Akçay & Beecher, 2015; Goodwin & Podos, 2014, 2015; Kroodsma, 2017). In chipping sparrows, syllable rate in particular has been shown to correlate with displays of territory defense: “birds responded more vigorously when simulated intruders sang the more difficult to produce, faster songs, and also when there was a stronger disparity between intruder trill rates and their own” (Goodwin & Podos, 2014). Some evidence suggests that chipping sparrows are subject to a performance constraint, specifically one in which there is a tradeoff between large sweeps in frequency (Hz) and high rate of syllable delivery (Goodwin & Podos, 2014; Podos, 1997). It is proposed that physiological constraints contribute to this balance in song performance (Podos, 1996, 1997). Further studies have suggested that this performance tradeoff between frequency bandwidth and syllable rate could be meaningful: under the stress of competing with the noise of an urban environment, chipping sparrows under-performed, singing “twice as far below the trade-off frontier” than those in less-noisy environments (Davidson et al., 2017). Kroodsma presents a contrary view based on results from field studies that demonstrate juvenile chipping sparrows imitate their neighbors with great success, and suggests that physiological constraints do not inhibit juveniles from performing fast songs (Kroodsma, 2017). Instead, he suggests that their performance is determined by that of their neighbor.

Our analysis is a step towards resolving the debate between performance-driven and neighbor-dependent hypotheses. These results suggest that chipping sparrows *select* which of their neighbors will be a tutor based on some aspect of their song performance: certain tutors may be preferred for reasons other than the how frequently the song is heard locally. Our analysis of recorded songs and song-learning models points to directional selection as the best explanation for chipping sparrow song diversity. In nature, juvenile chipping sparrows sing several neighbors’ songs before selecting a final song, which suggests that a selective process is taking place (W.-C. Liu & Nottebohm, 2007). This selective process, along with juveniles’ modification of their song during the plastic phase of song learning, have been proposed to play a part in determining their final song (Podos, 2017). The extent to which these potential selective processes affect song learning is controversial, suggesting that the combination of song data with learning simulations could shed light on the evolutionary dynamics of vocal learning.

Our 70-year sampling timespan gave us the opportunity to analyze the observed longevity of chipping sparrow syllable types alongside their frequency of occurrence. We find that it is difficult to reproduce the distribution of syllable occurrences in our regional analyses (**Figs. 4, 5**), but the distribution of syllable lifespans was only reproduced by models of directional selection. This divergence seems to be driven by the models predicting a large number of uncommon, short-lived syllables. Overall, these spectra of syllable properties favor the directional model of song transmission in chipping sparrows, such that some quality of the song is under selection, rather than its frequency in the local population. Lachlan *et al*. demonstrated a model of conformity bias in swamp sparrows that leads to a qualitatively similar lifespan distribution as ours, in which certain syllables tend to be longer-lived, even predicting that these syllables are maintained for upwards of 500 years (Lachlan et al., 2018). In contrast to their model of swamp sparrows, our chipping sparrow model supports directional selection as the more likely source of the observed patterns of syllable lifespan (**Figs. 4-5** for the NY region, **Figs. S4-5** for all others). A major difference between our model and that of Lachlan et al. is that we explicitly modeled the spatial structure of songbird populations, such that conformity bias only acts on the syllables found among neighbors. As a result, the conformity factor that we find to be most appropriate (*c*=1.2) cannot be directly compared to the parameter α used byLLachlan et al., which they find fits their swamp sparrow data best at 𝛼=1.316.

We compared the results of our model to empirical data from three focal regions—each having a high density of song recording coverage—and we found the same patterns applied to all of these regions. Directional selection produced the best result in all three regions, consistently favoring low learning error rates (<0.1% error) and some amount of dispersal (dispersal rate ≥0.1) (for a heatmap of the NY region, see **Fig. 4**; for OH/MI and New England, see **Fig. S6** panels G-R; for best-fit results for all regions, see **Table S4**). The comparison of the model to the entire range produced different results: in this case, conformity biased learning can also reproduce the empirical distribution of syllable lifetimes (**Fig. S6** panel E). The directional model of selection consistently produced the syllable lifetimes found in all regions, including a number of long-lived syllables. However, the directional learning strategy never produced a good fit for the empirical frequencies of syllable occurrence for a matrix size of 500×500. Even the best-fit models tended to underestimate the number of very common syllable types (**Figs. 5, S4**). Stronger selective pressures may cause syllables to be more common in these models, leading to better estimates of syllable occurrences and lifespans.

Several reasons for the differences between the model results and empirical data are suggested by the patterns in our results. We find that smaller models of directional selection (with 160,000 territories) effectively describe the empirical distributions of song occurrences and lifetimes for a wide range of parameters (**Figs. S15**), whereas models with population sizes closer to our estimates (up to 490,000 territories, compared to 250,000 in most of our models and from 444,000 to 833,000 in these regions) are less effective (**Figs. S13-15, Table S5**). A major factor that can explain this discrepancy is the difference in sampling: the community-science samples are not randomly distributed, whereas those of our model are. Song recordings are most common at the intersection of high human and high chipping sparrow population densities. This sampling discrepancy could mean that the empirical samples capture a much smaller effective population of chipping sparrows than exists in the entire region. In addition, the range of song rates (∼36.5–40 syllables per second) observed in the entire simulated population (before sampling) is much higher and narrower than that observed in the chipping sparrow population (∼5–38 syllables per second) (Searfoss, Liu, et al., 2020). This supports our intuition that syllable rate is not under directional selection on its own, since we previously observed a wide range of syllable rates in chipping sparrow songs that persisted over many years in nature (Searfoss, Liu, et al., 2020).

This model does not reproduce the entire song learning process and, since there has been a single detailed study on the chipping sparrow song-learning process (W.-C. Liu & Kroodsma, 2006), we do not know whether they learn identically across their range. However, our results suggest that chipping sparrows learn songs with a preference for one or several song features in at least part of their range. The presence of significant local diversity (W.-C. Liu & Kroodsma, 2006; Marler & Isaac, 1960) and distribution of multiple syllables across the country and overlapping in the same region (Searfoss, Liu, et al., 2020) also suggests that chipping sparrows do not have a strong conformist bias in their learning. Our results can be compared to those in house finches, which demonstrated content bias—certain syllables were more likely to be learned because of their acoustic features, not because of their frequency of occurrence in the population (Youngblood & Lahti, 2022). In chipping sparrows, since only one syllable is learned per bird, we tracked potential selection on the acoustic features themselves to test whether directional selection favoring the learning of faster songs can explain the observed distribution of syllables.

Sparrow species such as the white-throated sparrows sing in their wintering grounds, allowing for rapid transmission of birdsong after these birds migrate north (Otter et al., 2020). We did not include this effect, since all of our songs from breeding ranges were recorded outside of the winter months (Searfoss, Liu, et al., 2020), and chipping sparrows are not known to sing regularly during winter months (W. C. Liu & Kroodsma, 1999). Song learning in the wintering grounds may explain some of the observed song variation, including songs that are widely dispersed (**Fig. S16**), as birds may have more tutors to learn from. This additional learning step may homogenize the population or increase the strength of a conformity bias (causing common songs to become more common among birds sharing wintering grounds).

The divergences between our model and community-science data suggests that more complex evolutionary pressures or cultural transmission biases might be at play, such as performance tradeoffs or differing selection pressures for tutor selection compared to mate choice, which could be integrated into the model for future research. One such explanation is a hypothesized performance tradeoff in chipping sparrow song between frequency bandwidth and the rate of syllable delivery (Podos, 1996, 1997); due to physiological constraints, a high-performance song might have a large frequency bandwidth but slower syllable rate or a faster syllable rate but reduced frequency bandwidth. In this case, directional selection is likely occurring on multiple axes and operating on both traits at once. If there is a tradeoff between two song parameters under selection, we would not expect to see tight distributions of a single syllable parameter (syllable rate or frequency modulation) as there will be a boundary along which the properties are balanced. Furthermore, although long-lived syllables had significantly shorter durations overall, we find a wide distribution of mean syllable durations, implying that both long and short syllables can persist over time.

It is difficult to determine whether a certain song feature is being favored by directional selection without corresponding field experiments. We framed directional tutor selection in our model such that a parameter was the determining factor for the learned syllable type. As a result, our simulations only suggest that a song characteristic could be under directional selection, not that syllable rate in particular is under selection. To test which song features might be favored in learning and tutor selection, we propose playback experiments to determine whether there is a difference in juveniles’ responses to recordings of different song rates, frequency bandwidths, and syllable complexities as well as to historical versus current song recordings (as in (Derryberry, 2007)). These results could then be compared to females’ responses to determine whether tutor selection and mate choice are favoring similar song characteristics. Ideally, this would be carried out at multiple locations across the chipping sparrow’s range, given the geographical patterns observed earlier. We aim to extend this model to incorporate content bias more broadly, allowing selection on the syllables themselves rather than aspects of syllable production such as syllable rate, as this may better align with empirical data in which buzz syllable types are long-lived. To execute this extension to the model, it would be necessary to create a measure of syllable quality to drive tutor choice.

We demonstrate that coupling an agent-based model with analyses of community-science data is a tool to better understand the evolution of behavior in a songbird. By developing a model of cultural transmission of song and comparing patterns produced by three learning strategies to those found in our empirical data, we demonstrate that the observed distributions of chipping sparrow syllable types show evidence of transmission bias. In particular, our results are indicative of a song-learning strategy in which tutor selection is under directional selection pressure in the chipping sparrow population, with juveniles preferentially selecting tutors with certain song features, and in which copy-errors or invention rates are quite low (<0.1%). While our simulation does not specify which specific features of syllables or songs are under selection for, we find that neutral song-learning processes and conformity-biased learning, both of which have been observed in other species (Lachlan et al., 2018; Slater, 1986), cannot explain the distribution of songs observed in chipping sparrows. Despite their deceptively simple song, our computational analyses suggest that chipping sparrows appear to be exhibiting learning biases and complex cultural transmission patterns, warranting further investigation in the field.

## Acknowledgements

We are indebted to the contributors and maintainers of the following databases of birdsong: Xeno-canto, Macaulay Library at the Cornell lab of Ornithology, and Borror Laboratory of Bioacoustics. We are grateful to Wan-chun Liu for his previous studies on chipping sparrows and the contribution of his field recordings of chipping sparrow songs. We thank the members of the Creanza Laboratory for their feedback. Y.P and N.C. were supported by the John Templeton Foundation (62187). A.M.S was supported by the National Science Foundation Graduate Research Fellowship Program under Grant DGE-1445197, and all authors received support from Vanderbilt University.

## Author Contributions

N.C. and A.M.S conceived and designed the project with input from Y.P. Y.P and A.M.S. coded the model and conducted all analyses with assistance from N.C. Y.P. and A.M.S and N.C. wrote the manuscript.

## Competing Interest Statement

The authors declare no competing interests.

## Data and materials availability

All code is available at https://github.com/CreanzaLab/ChippingSparrowCulturalEvolutionModel, For the catalog numbers, database, recordist, URL, and license for the 820 song files, see **Supplementary Data Table 1**. For the metadata including recording latitudes and longitudes and the 8 song features (all log transformed except mean stereotypy of repeated syllables and the standard deviation of note frequency modulation), see **Supplementary Data Table 2**.

## Notes

https://github.com/CreanzaLab/ChippingSparrowCulturalEvolutionModel

### Competing Interest Statement

The authors have declared no competing interest.

### Summary of Updates

This version of the manuscript has an expanded set of regional simulations; we tested a broader parameter range and explored how the size of our simulations is associated with the potential number of chipping sparrow territories in each of our focal regions.

