## Supplemental Tables and Figures for "Detecting cultural evolution in a songbird species using community-science data and computational modeling"

**Supplementary Data Table S1** Catalog numbers, database, recordist, URL, and license for the 820 song files. See attached file.

**Supplementary Data Table S2** Metadata for the 820 song files, including recording latitudes and longitudes and the 8 song features (all log transformed except mean stereotypy of repeated syllables and the standard deviation of note frequency modulation). See attached file.

**Supplementary Data Table S3** Best parameters for different models in the NY region. *P*-values representing simulated data statistically indistinguishable from the empirical data in bold. For best parameters of other regions, see **Table S4**.

| Model type | Best parameters | | Counts *P*-value | Lifetimes *P*-value |
|---|---|---|---|---|
| | learning error rate | dispersal fraction | | |
| Neutral | $10^{-5}$ | 0.3 | $5.63\times10^{-13}$ | $9.46\times10^{-6}$ |
| Conformity (*c*=1.2) | 0.001 | 0.1 | $4.56\times10^{-8}$ | $8.20\times10^{-4}$ |
| Conformity (*c*=1.4) | 0.01 | 0.6 | $3.22\times10^{-11}$ | $1.48\times10^{-4}$ |
| Conformity (*c*=1.6) | 0.1 | 0.9 | $6.53\times10^{-13}$ | $6.02\times10^{-5}$ |
| Conformity (*c*=1.8) | $10^{-5}$ | 0.3 | $3.44\times10^{-13}$ | $3.15\times10^{-5}$ |
| Conformity (*c*=2) | 0.01 | 0.4 | $3.44\times10^{-13}$ | $3.15\times10^{-5}$ |
| Novelty (*c*=0.8) | $10^{-5}$ | 0.1 | $1.1\times10^{-15}$ | $1.09\times10^{-6}$ |
| Directional | $10^{-6}$ | 0.5 | 0.00441 | **0.0123** |

**Supplementary Data Table S4** Best parameters for different models and regions. All models are for a matrix size of 500×500. *P*-values representing simulated data statistically indistinguishable from the empirical data in bold.

| Model type | Region | Best parameters | | Counts *P*-value | Lifetimes *P*-value |
|---|---|---|---|---|---|
| | | Learning error rate | Dispersal fraction | | |
| Neutral | Entire range | 0.001 | 1.0 | $1.76\times10^{-42}$ | $9.25\times10^{-4}$ |
| | NY | $10^{-5}$ | 0.3 | $5.63\times10^{-13}$ | $9.46\times10^{-6}$ |
| | OH/MI | 0.0001 | 0.1 | $6.34\times10^{-22}$ | $3.16\times10^{-8}$ |
| | New England | 0.0001 | 0.6 | $4.71\times10^{-28}$ | $3.63\times10^{-9}$ |
| Conformity (*c*=1.2) | Entire range | 0.001 | 0.1 | $6.91\times10^{-17}$ | **0.109** |
| | NY | 0.01 | 0.4 | $3.44\times10^{-13}$ | $3.15\times10^{-5}$ |
| | OH/MI | 0.001 | 0.1 | $1.07\times10^{-10}$ | $1.35\times10^{-4}$ |
| | New England | 0.001 | 0.1 | $5.82\times10^{-13}$ | $3.63\times10^{-5}$ |
| Directional | Entire range | 0.01 | 0.5 | $3.08\times10^{-4}$ | **0.136** |
| | NY | $10^{-6}$ | 0.5 | 0.00441 | **0.123** |
| | OH/MI | $10^{-6}$ | 0.5 | 0.00367 | **0.0762** |
| | New England | 0.0001 | 0.5 | $1.41\times10^{-4}$ | **0.0545** |

**Supplementary Data Table S5** Best parameters for different models and different matrix sizes for the NY region. *P*-values representing simulated data statistically indistinguishable from the empirical data in bold.

| Model type | Matrix dimension | Best parameters | | Counts *P*-value | Lifetimes *P*-value |
|---|---|---|---|---|---|
| | | Learning error rate | Dispersal fraction | | |
| Neutral | 400 | 0.0001 | 1.0 | $1.62\times10^{-12}$ | $1.73\times10^{-4}$ |
| | 500 | $10^{-5}$ | 0.3 | $5.63\times10^{-13}$ | $9.46\times10^{-6}$ |
| | 600 | 0.001 | 0.6 | $1.84\times10^{-14}$ | $2.50\times10^{-6}$ |
| | 700 | $10^{-5}$ | 0.8 | $2.15\times10^{-15}$ | $8.32\times10^{-7}$ |
| Conformity (*c*=2) | 400 | 0.001 | 0.7 | $1.67\times10^{-10}$ | $4.36\times10^{4}$ |
| | 500 | 0.01 | 0.4 | $3.44\times10^{-13}$ | $3.15\times10^{-5}$ |
| | 600 | $10^{-5}$ | 0.2 | $3.37\times10^{-14}$ | $5.22\times10^{-6}$ |
| | 700 | 0.001 | 0.6 | $1.82\times10^{-15}$ | $6.15\times10^{-7}$ |
| Directional | 400 | $10^{-4}$ | 0.4 | **0.189** | **0.345** |
| | 500 | $10^{-6}$ | 0.5 | 0.00441 | **0.123** |
| | 600 | $10^{-5}$ | 0.5 | $1.73\times10^{-5}$ | 0.00871 |
| | 700 | 0.001 | 0.5 | $7.90\times10^{-9}$ | 0.00222 |

**Supplementary Data Table S6** Best parameters for different models of syllable learning error. In the standard model, errors result in novel syllable invention; in the model with homoplasy, syllables are chosen from a fixed set of 500 syllables, identical with the syllables from the model initialization. All values represent those for the NY region with a matrix size of 500×500. *P*-values representing simulated data statistically indistinguishable from the empirical data in bold.

| Model type | Homoplasy | Best parameters | | Counts *P*-value | Lifetimes *P*-value |
|---|---|---|---|---|---|
| | | Learning error rate | Dispersal fraction | | |
| Neutral | No | $10^{-5}$ | 0.3 | $5.63\times10^{-13}$ | $9.46\times10^{-6}$ |
| | Yes | 0.0001 | 0.6 | $8.58\times10^{-14}$ | $1.134\times10^{-5}$ |
| Conformity (*c*=2) | No | 0.01 | 0.4 | $3.44\times10^{-13}$ | $3.15\times10^{-5}$ |
| | Yes | $10^{-5}$ | 0.7 | $1.20\times10^{-14}$ | $2.31\times10^{-5}$ |
| Directional | No | $10^{-6}$ | 0.5 | 0.00441 | **0.123** |
| | Yes | 0.01 | 1.0 | $0.00270^{3}$ | **0.0809** |

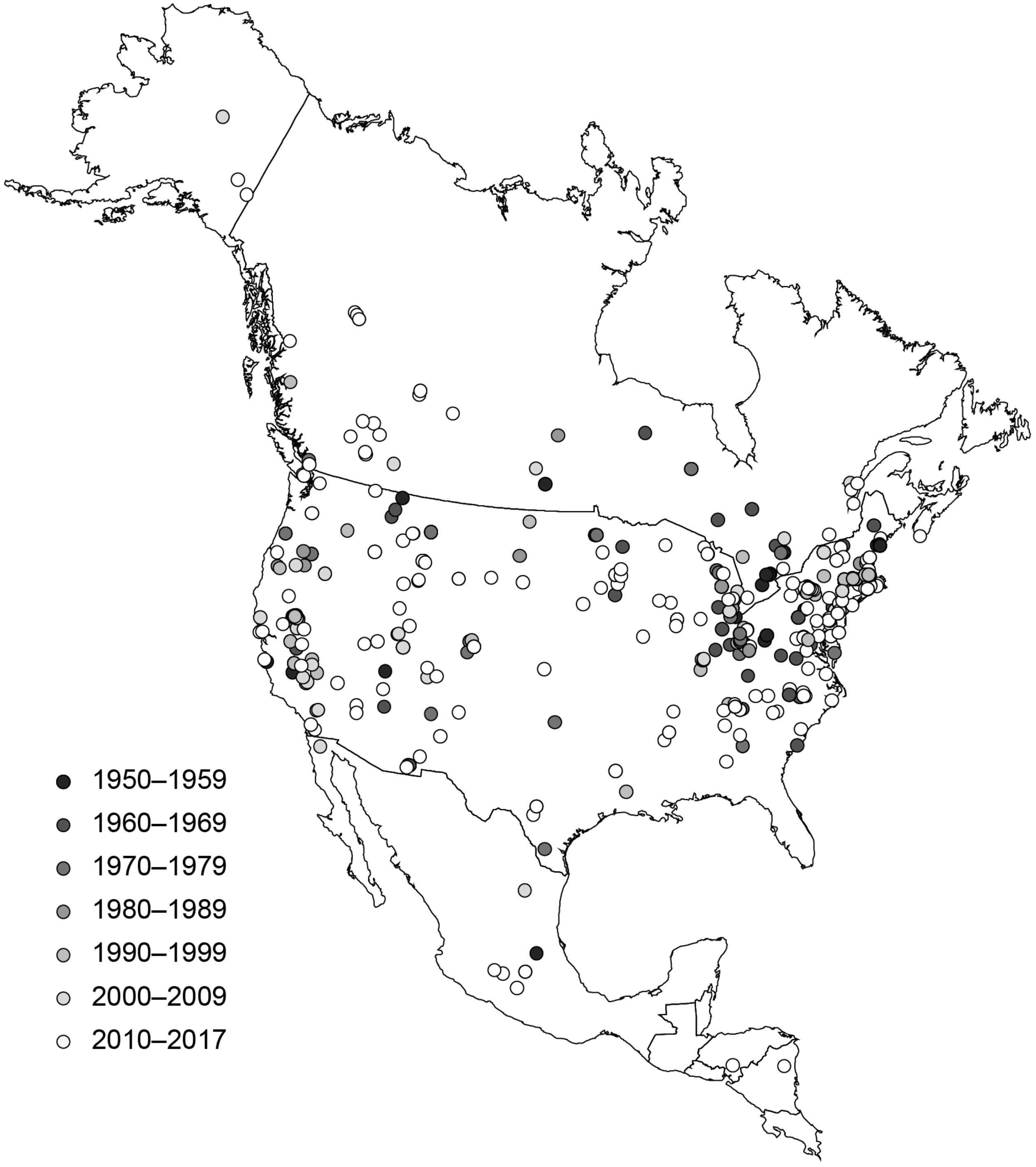


**Supplementary Figure S1. Locations and decades of the 820 collected chipping sparrow recordings.** The Geographic map was made using ArcMap v.10.7 and country outlines are from Esri (DeLorme Publishing Company, Inc., map projection, North_America_Lambert_Conformal_Conic, WKID: 102009 Authority: Esri).

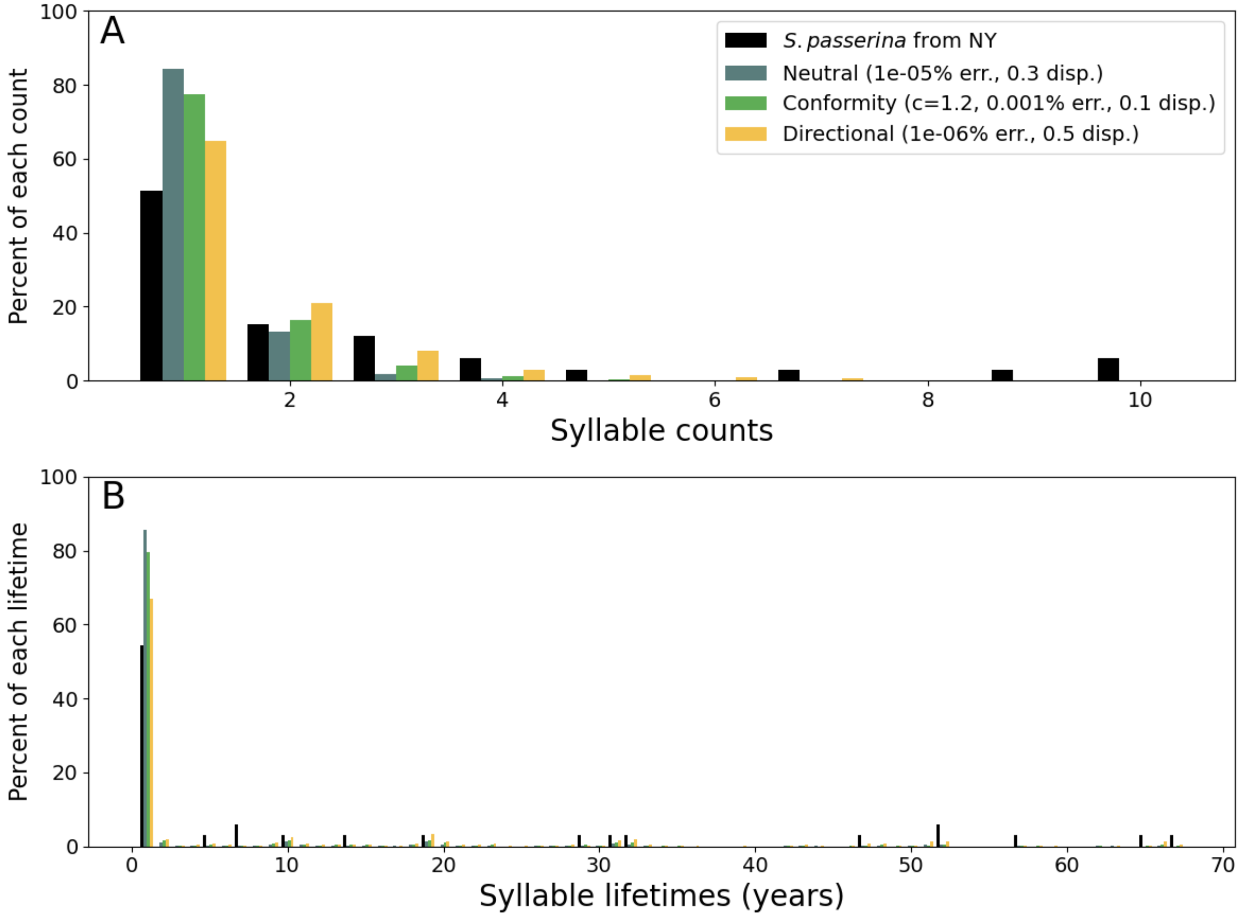


**Supplementary Figure S2. Unbinned data from Fig. 5. Unbinned syllable syllable frequency spectra and syllable lifespan spectra of empirical data and of samples from best-fit models.** (A) The percent of syllable types versus the frequency at which the syllable types occur in the population (e.g. the number of birds sampled with that syllable type) for the best-fit models of each of the three song-learning strategies: (blue) neutral tutor selection, (green) conformity bias (*c*=1.2), and (yellow) directional selection. In each panel, the histogram from community-science data is also provided (black). (B) The percent of syllable types versus the syllable lifespans (calculated as the last year − the first year in which the syllable type was sampled) for the best-fit models of each of the song learning strategies. For simulated data, 50 resamplings were taken to estimate likely See also **Fig. S3**.

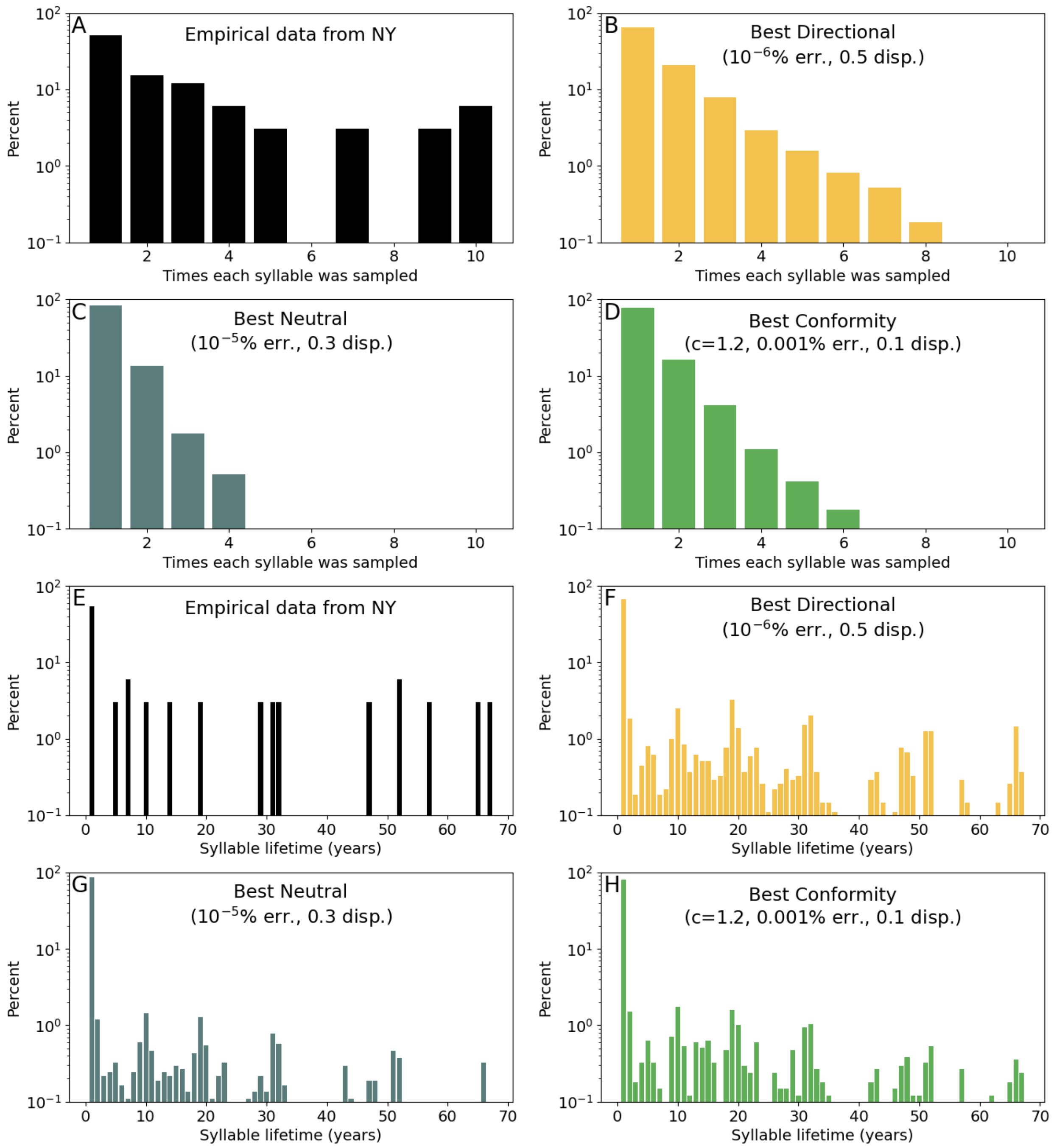


**Supplementary Figure S3. Unbinned, log-scaled plot of the data found in Fig. 5. Unbinned syllable frequency spectra (A-D) and syllable lifespan spectra (E-H) of empirical data from the NY region and samples from best-fit models, split for legibility.** For visualization, these plots have a log-scaled y-axis. See **Fig. S2** for details.

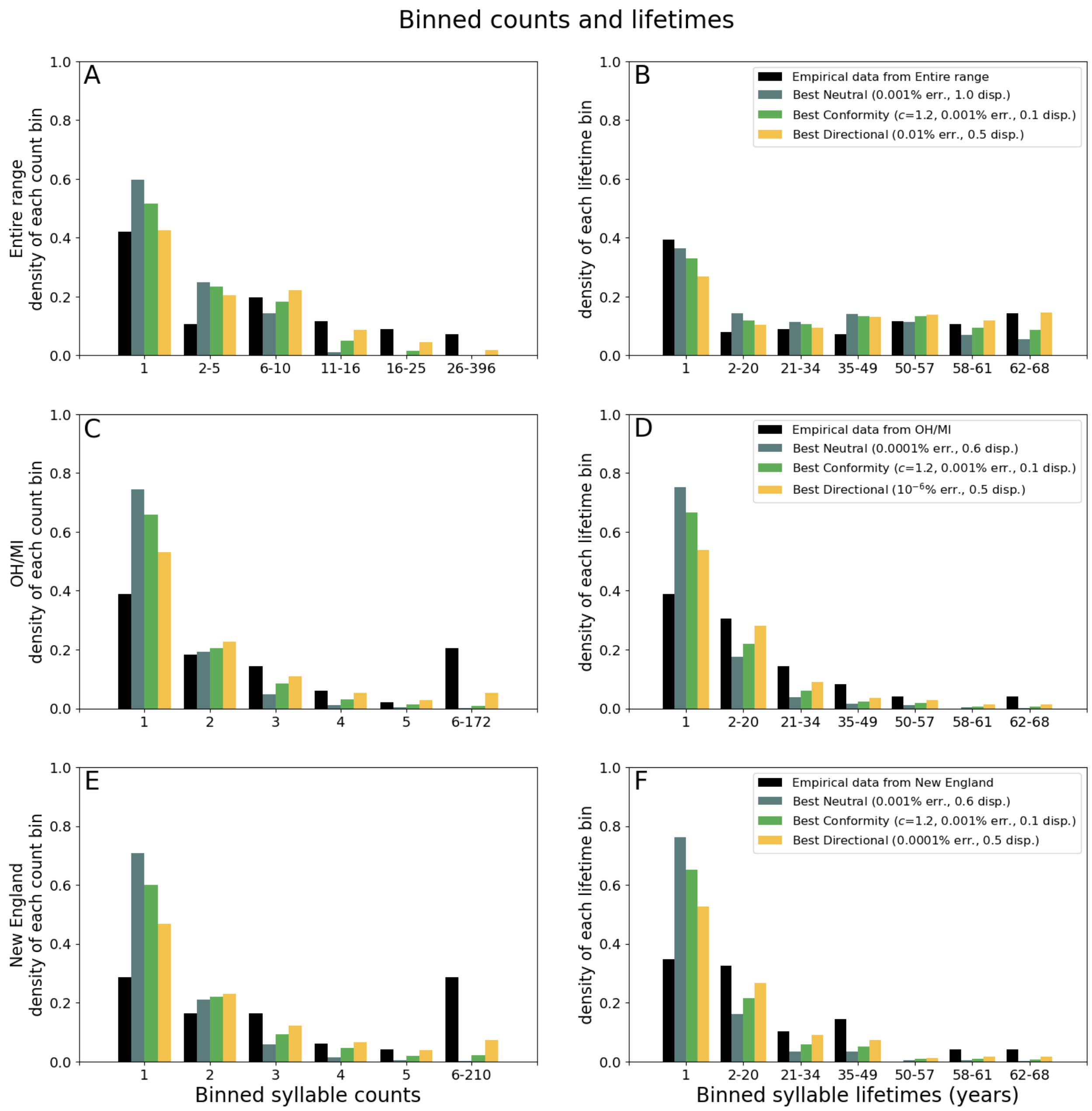


**Supplementary Figure S4. Similar to Fig. 5, but for different regions. Binned syllable frequency spectra and syllable lifespan spectra from focal regions and samples from best-fit models.** (A,C,E) The percent of syllable types versus the frequency at which the syllable types occur in the population (e.g. the number of birds sampled with that syllable type) for the best-fit models of each of the three song-learning strategies. In each panel, the histogram from community-science data is also provided (black). (B,D,F) The percent of syllable types versus the syllable lifespans (calculated as the last year − the first year in which the syllable type was sampled) for the best-fit models of each of the song learning strategies. See also **Fig. S5** for unbinned data.

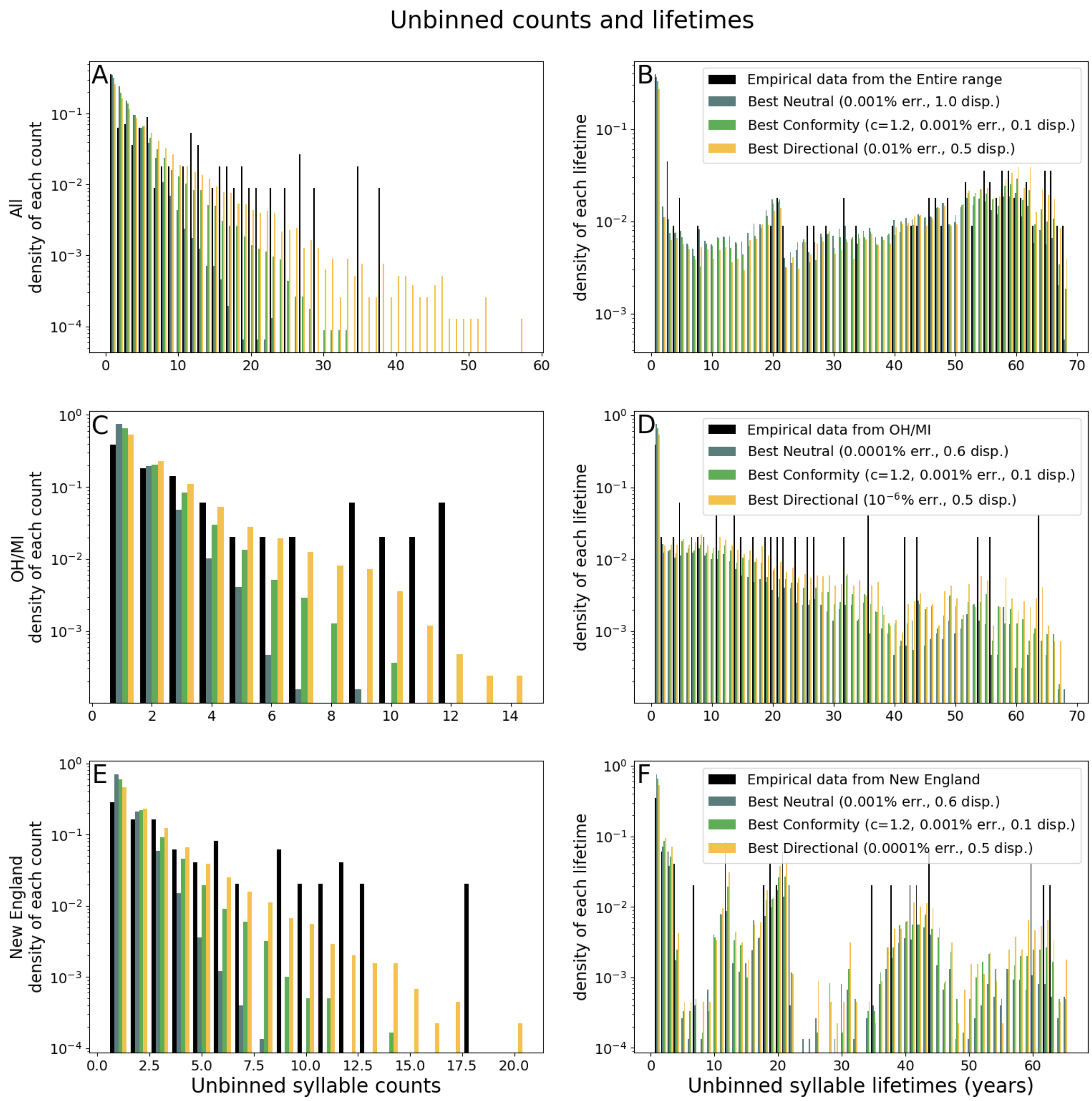


**Supplementary Figure S5. Similar to Fig. 5, but for different regions and with log-scaled and unbinned data. Unbinned syllable frequency spectra and syllable lifespan spectra from focal regions and samples from best-fit models.** For visualization, these plots have a log-scaled y-axis. See **Fig. S4**.

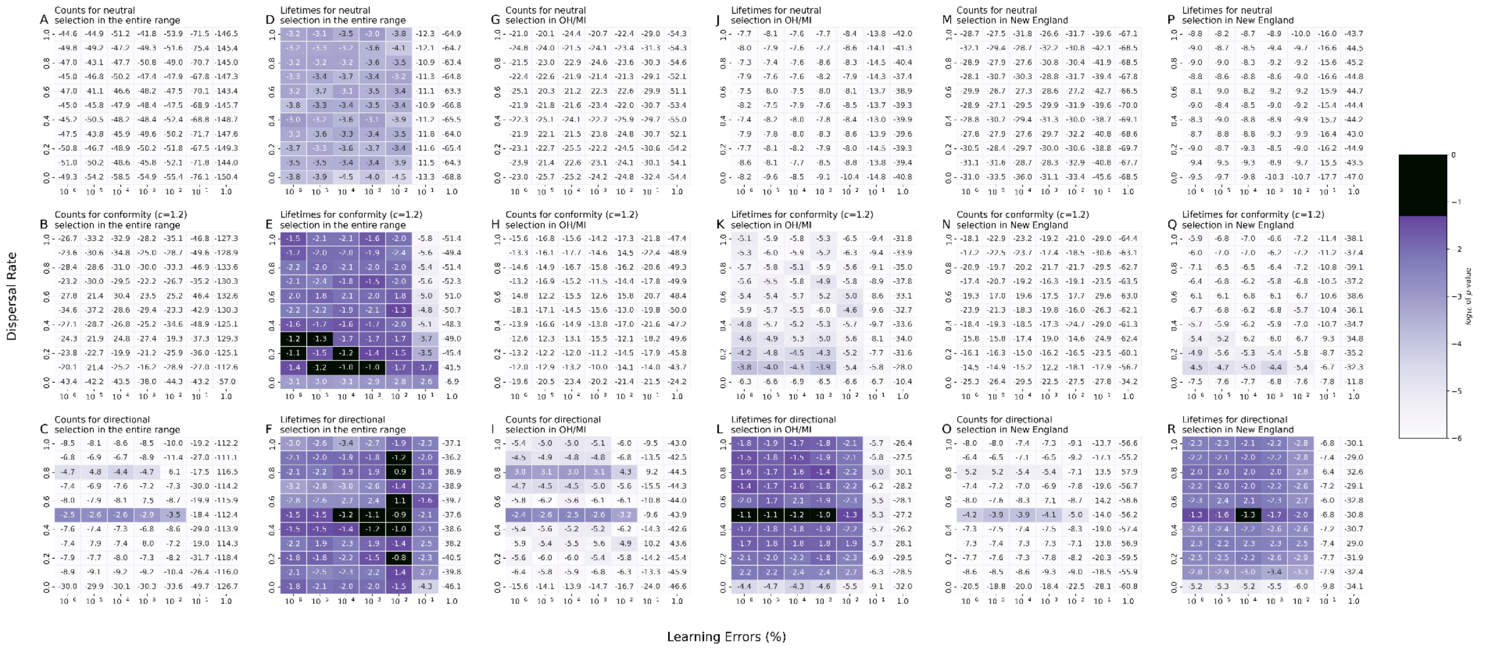

**Supplementary Figure S6. Similar to Fig. 4, but for the entire range (A-F), OH/MI (G-L), and New England (M-R). Comparisons of syllable count and lifespan frequency spectra between models of neutral, conformity, and directional selection with empirical data.** *P*-values for Fisher's exact tests in which the null hypothesis is that the binned syllable frequency spectra of empirical data and the simulated data are drawn from the same distribution. Models that are statistically indistinguishable (*P*>0.05) from the empirical data are shown in black.

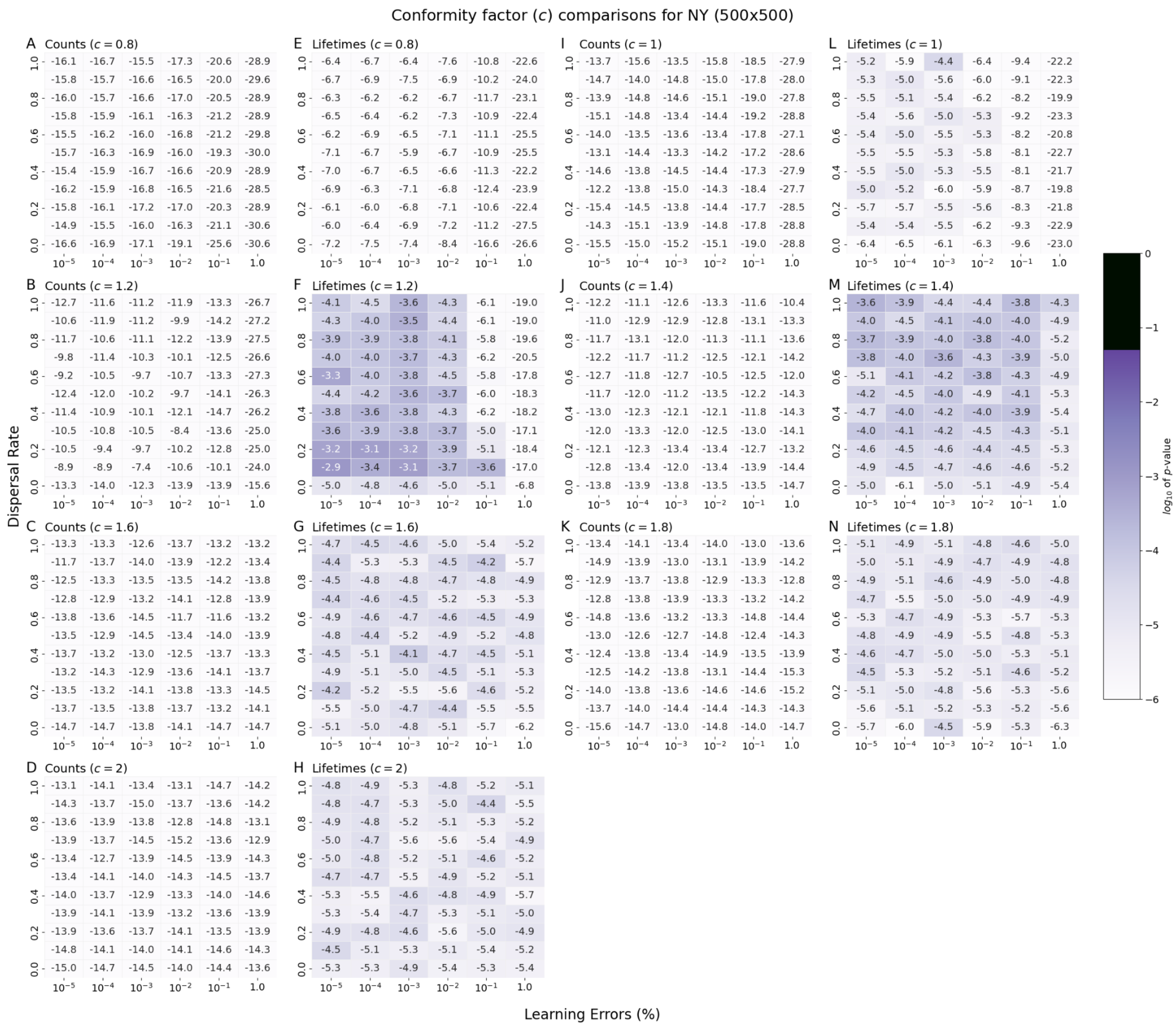


**Supplementary Figure S7. Comparisons of syllable count (A-D & I-K) and lifespan frequency spectra (E-H & L-N) between models with different conformity factors (*c* from 0.8 to 2.0 in increments of 0.2) and empirical data across a range of dispersal fractions and learning-error rates.** *P*-values for Fisher's exact tests in which the null hypothesis is that the binned syllable frequency spectra of empirical data and the simulated data are drawn from the same distribution. All simulations produced results that were significantly different from the empirical data ($P$>0.05).

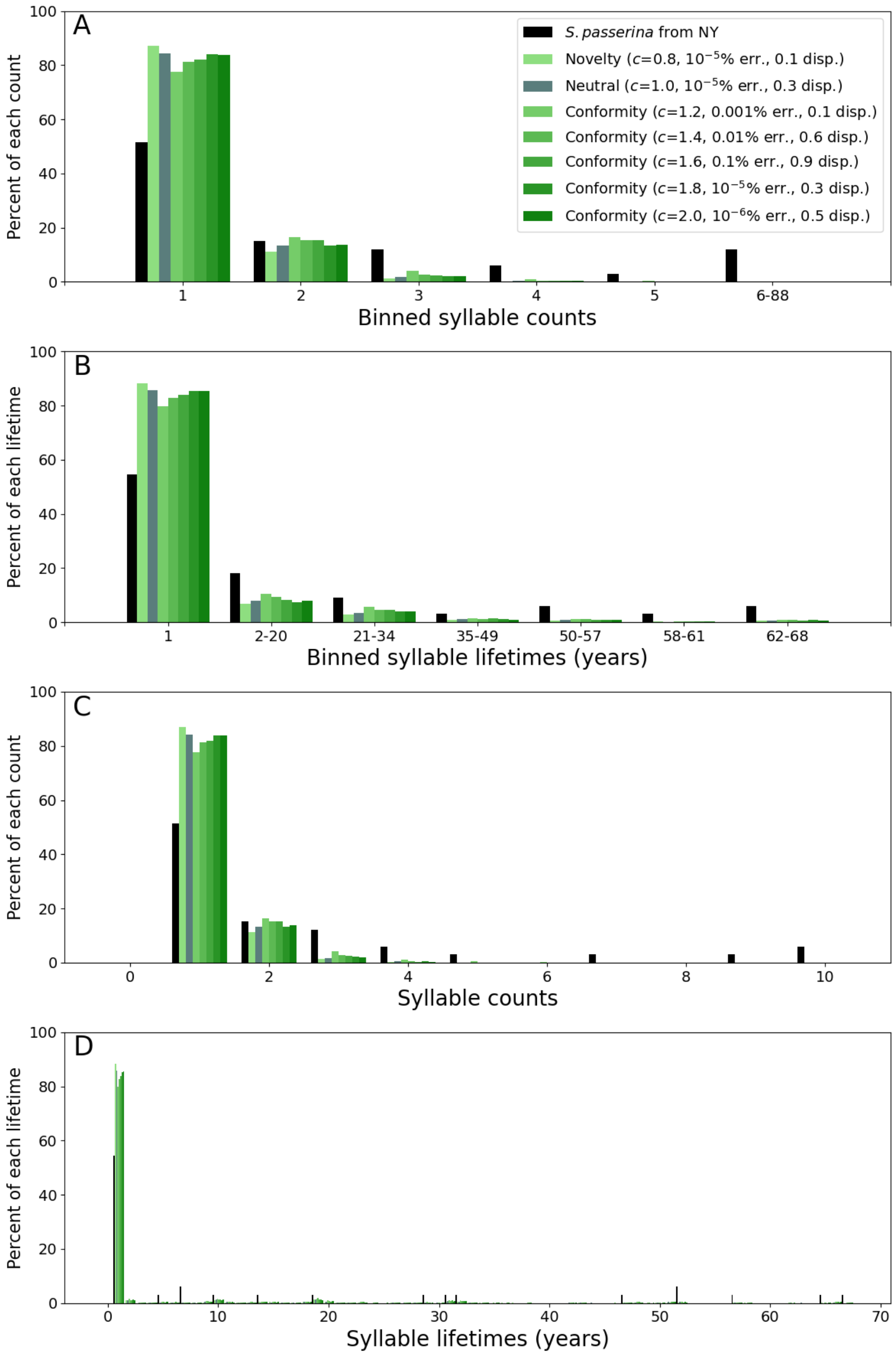


**Supplementary Figure S8. Syllable frequency and syllable lifespan spectra from empirical data (in black) and samples from the best-fit conformity models (in green and blue).** (A & C) The percent of syllable types versus the frequency at which the syllable types occur in the population (e.g. the number of birds sampled with the syllable type) for the best-fit models of each of the conformity models, including novelty (conformity factor $c$=0.8), neutral biased song transmission ($c$=1). In each panel, the histogram from citizen-science data is also provided (black). (B) The percent of syllable types versus the syllable lifespans for the best-fit models of each of the conformity models.

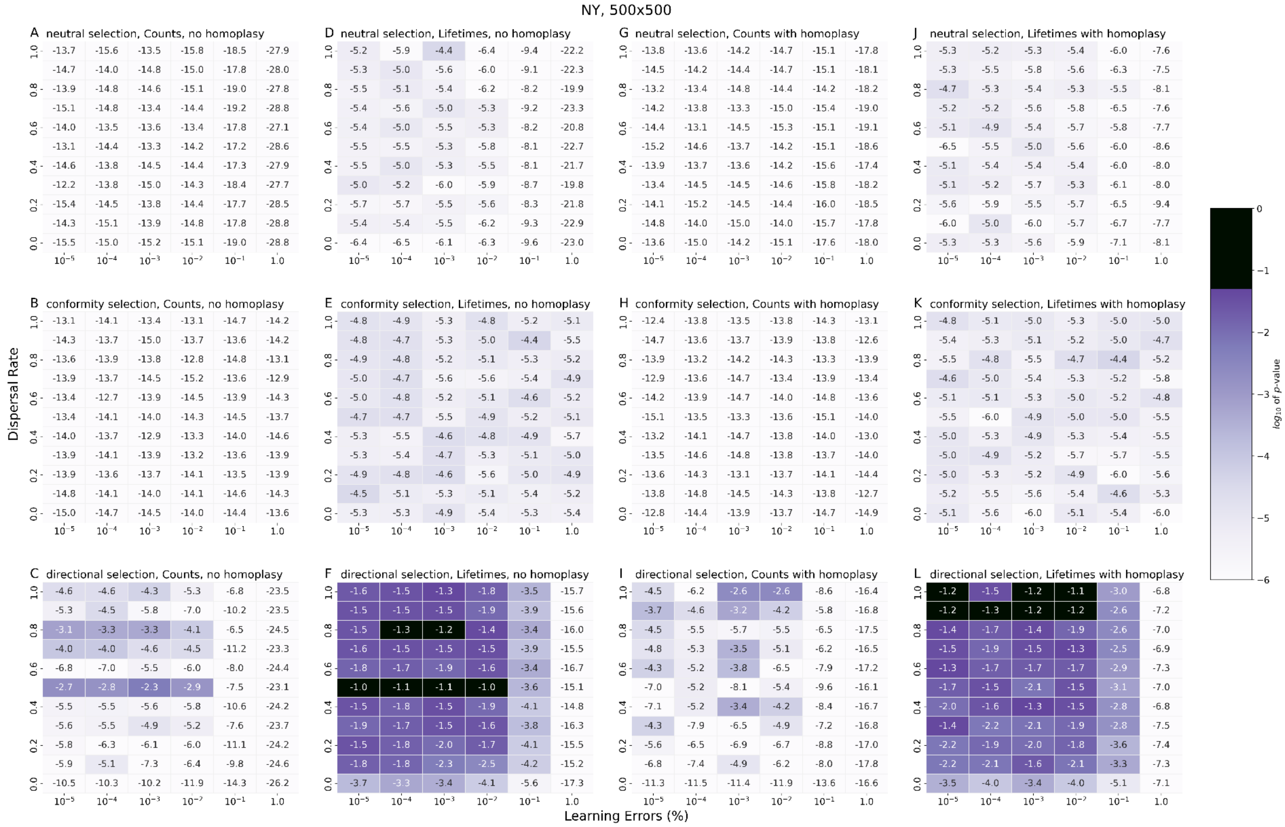


**Supplementary Figure S9. Comparisons of selection without (A-F) with homoplasy (G-L) for neutral, conformity, and directional selection. This includes counts (A-C & G-I) and frequency spectra (D-F & J-L) for each selection model compared to the NY region.** *P*-values for Fisher's exact tests in which the null hypothesis is that the binned syllable frequency spectra of empirical data and the simulated data are drawn from the same distribution. Models that are statistically indistinguishable ($P$>0.05) from the empirical data are shown in black. For other regions, see **Figs. S10-12**.

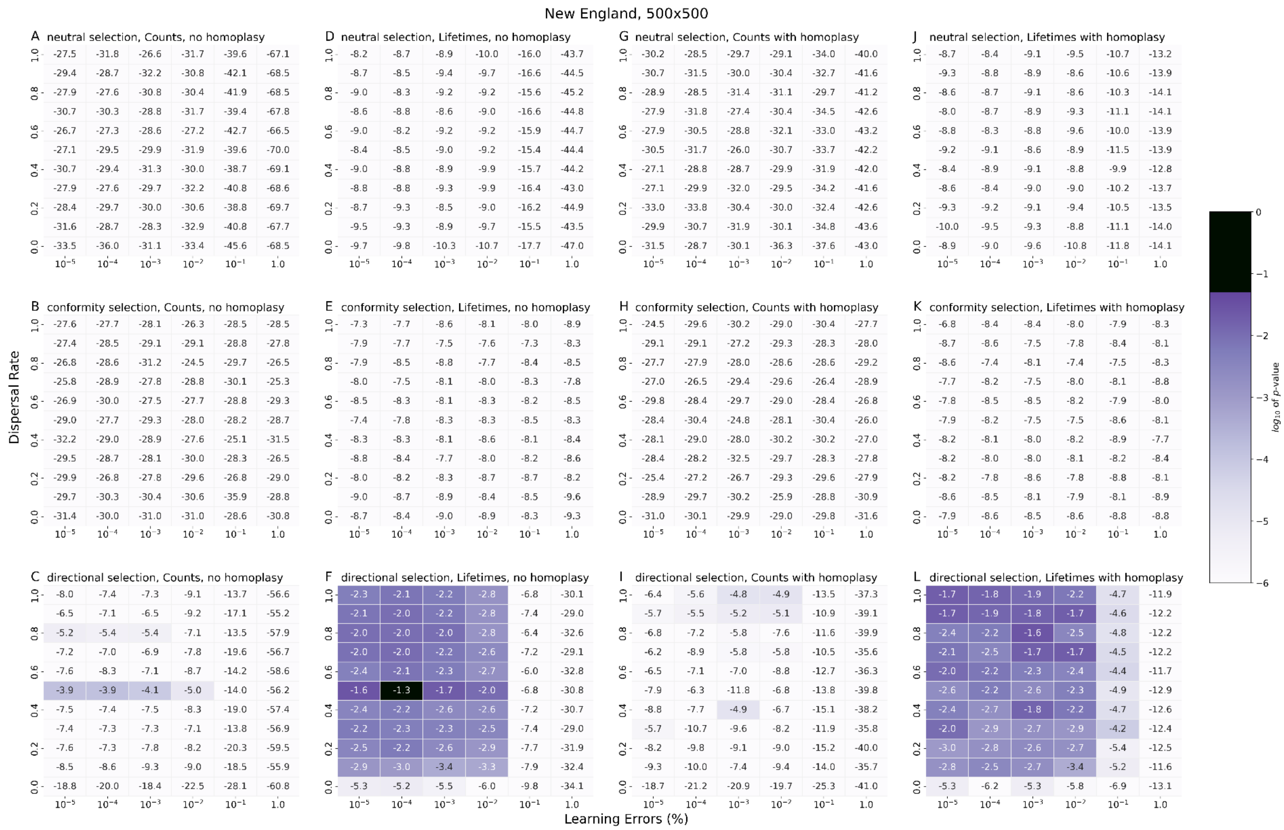


**Supplementary Figure S10. Similar to Fig. S8, but for the New England region. Comparisons of selection without (A-F) with homoplasy (G-L) for neutral, conformity, and directional selection. This includes counts (A-C & G-I) and frequency spectra (D-F & J-L) for each selection model compared to the New England region.** *P*-values for Fisher's exact tests in which the null hypothesis is that the binned syllable frequency spectra of empirical data and the simulated data are drawn from the same distribution. Models that are statistically indistinguishable (*P*>0.05) from the empirical data are shown in black. For other regions, see **Figs. S9, S11-12**.

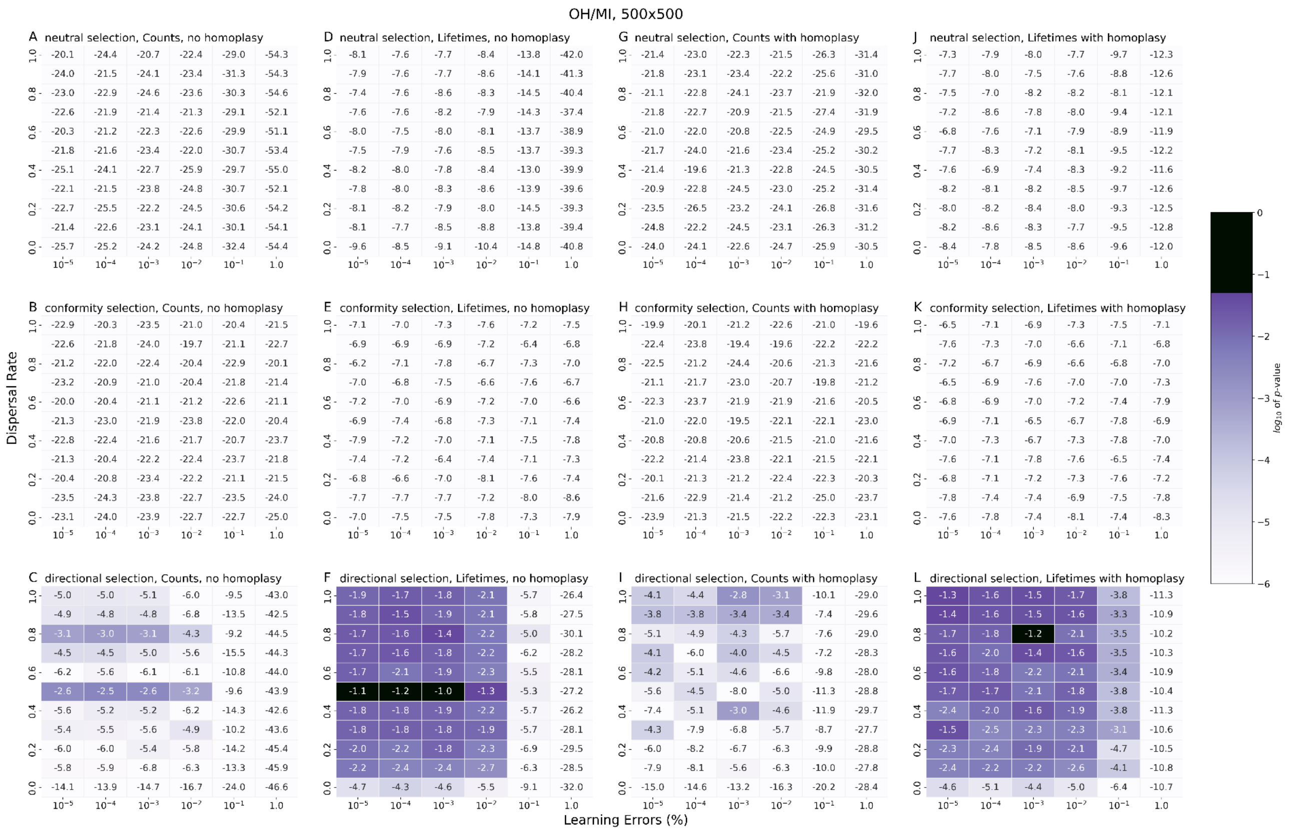


**Supplementary Figure S11. Similar to Fig. S8, but for the Ohio/Michigan region. Comparisons of selection without (A-F) with homoplasy (G-L) for neutral, conformity, and directional selection. This includes counts (A-C & G-I) and frequency spectra (D-F & J-L) for each selection model compared to the Ohio/Michigan region.** *P*-values for Fisher's exact tests in which the null hypothesis is that the binned syllable frequency spectra of empirical data and the simulated data are drawn from the same distribution. Models that are statistically indistinguishable (*P*>0.05) from the empirical data are shown in black. For other regions, see **Figs. S9-10, S12**.

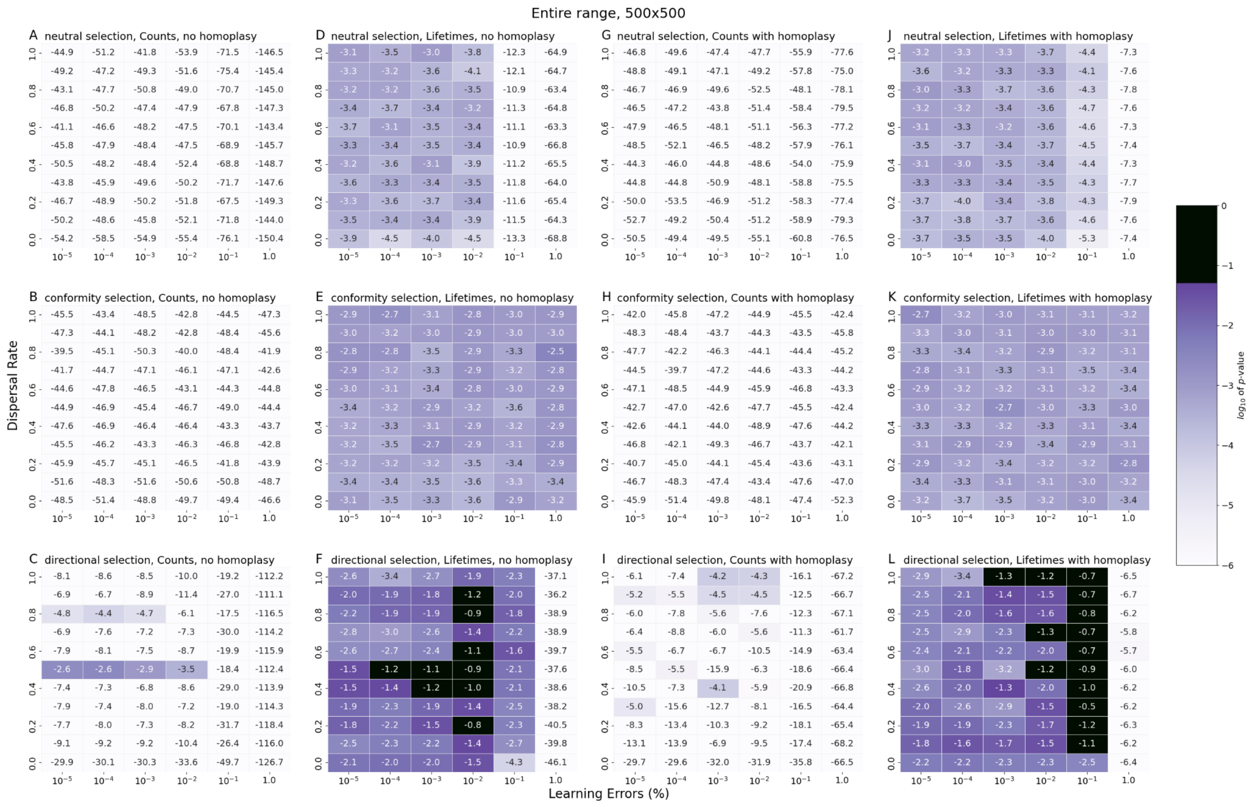

**Supplementary Figure S12. Similar to Fig. S8, but for the entire range. Comparisons of selection without (A-F) with homoplasy (G-L) for neutral, conformity, and directional selection. This includes counts (A-C & G-I) and frequency spectra (D-F & J-L) for each selection model compared to samples from the entire region.** *P*-values for Fisher's exact tests in which the null hypothesis is that the binned syllable frequency spectra of empirical data and the simulated data are drawn from the same distribution. Models that are statistically indistinguishable (*P*>0.05) from the empirical data are shown in black. For other regions, see **Figs. S9-11**.

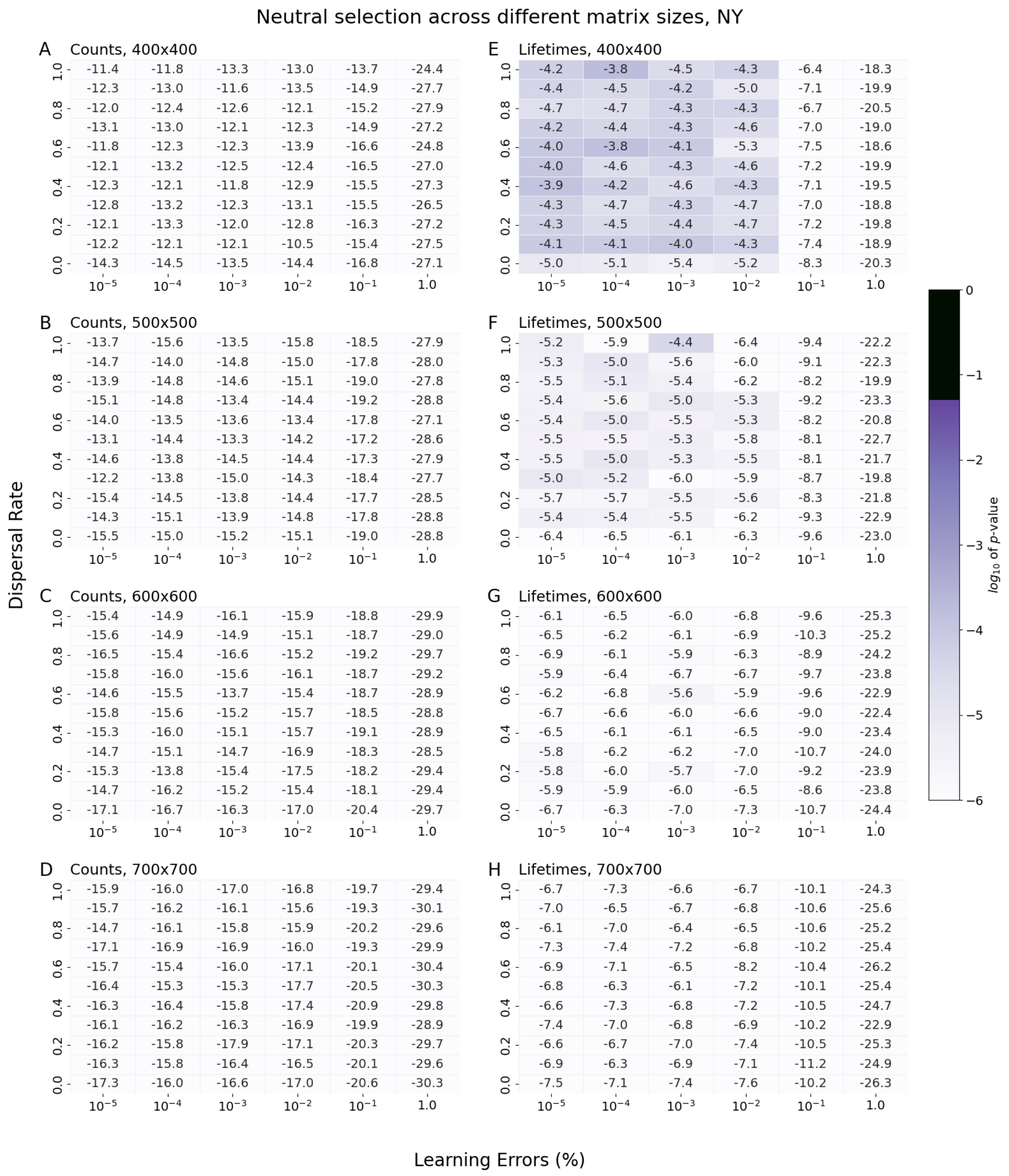


**Supplementary Figure S13. Comparisons of the effects of neutral selection across different matrix sizes (400 by 400 to 700 by 700 in increments of 100), including counts (A-D) and frequency spectra (E-H) compared to empirical data in the NY region.** *P*-values for Fisher's exact tests in which the null hypothesis is that the binned syllable frequency spectra of empirical data and the simulated data are drawn from the same distribution. All simulations produced results that were significantly different from the empirical data (P<0.05). For conformity and directional selection, see **Fig. S14-15**.

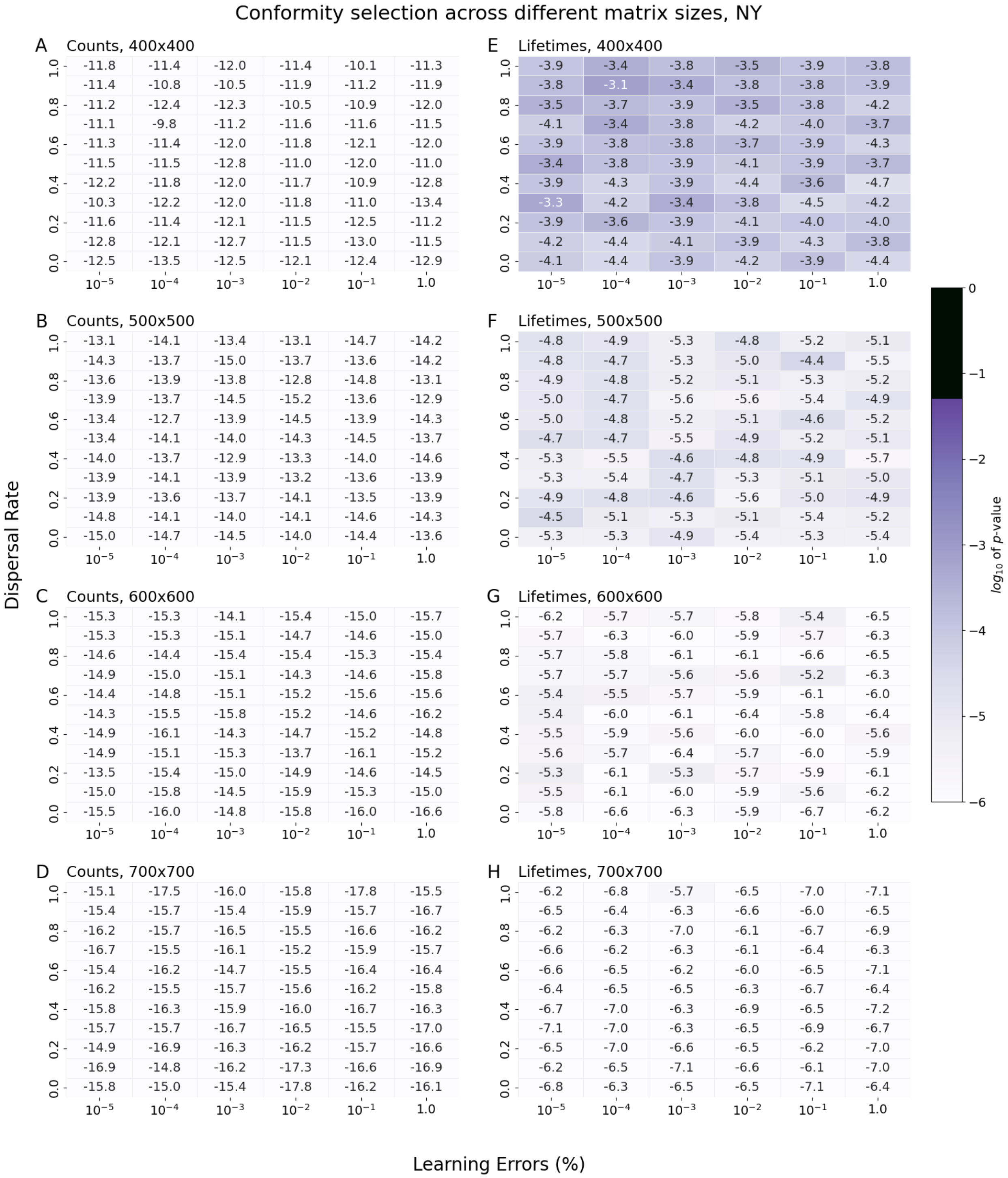


**Supplementary Figure S14. Comparisons of the effects of conformity selection across different matrix sizes (400 by 400 to 700 by 700 in increments of 100), including counts (A-D) and frequency spectra (E-H) compared to empirical data in the NY region.** *P*-values for Fisher's exact tests in which the null hypothesis is that the binned syllable frequency spectra of empirical data and the simulated data are drawn from the same distribution. All simulations produced results that were significantly different from the empirical data (P<0.05). For neutral bias, see **Fig. S13** and for directional selection, see **Fig. S15**.

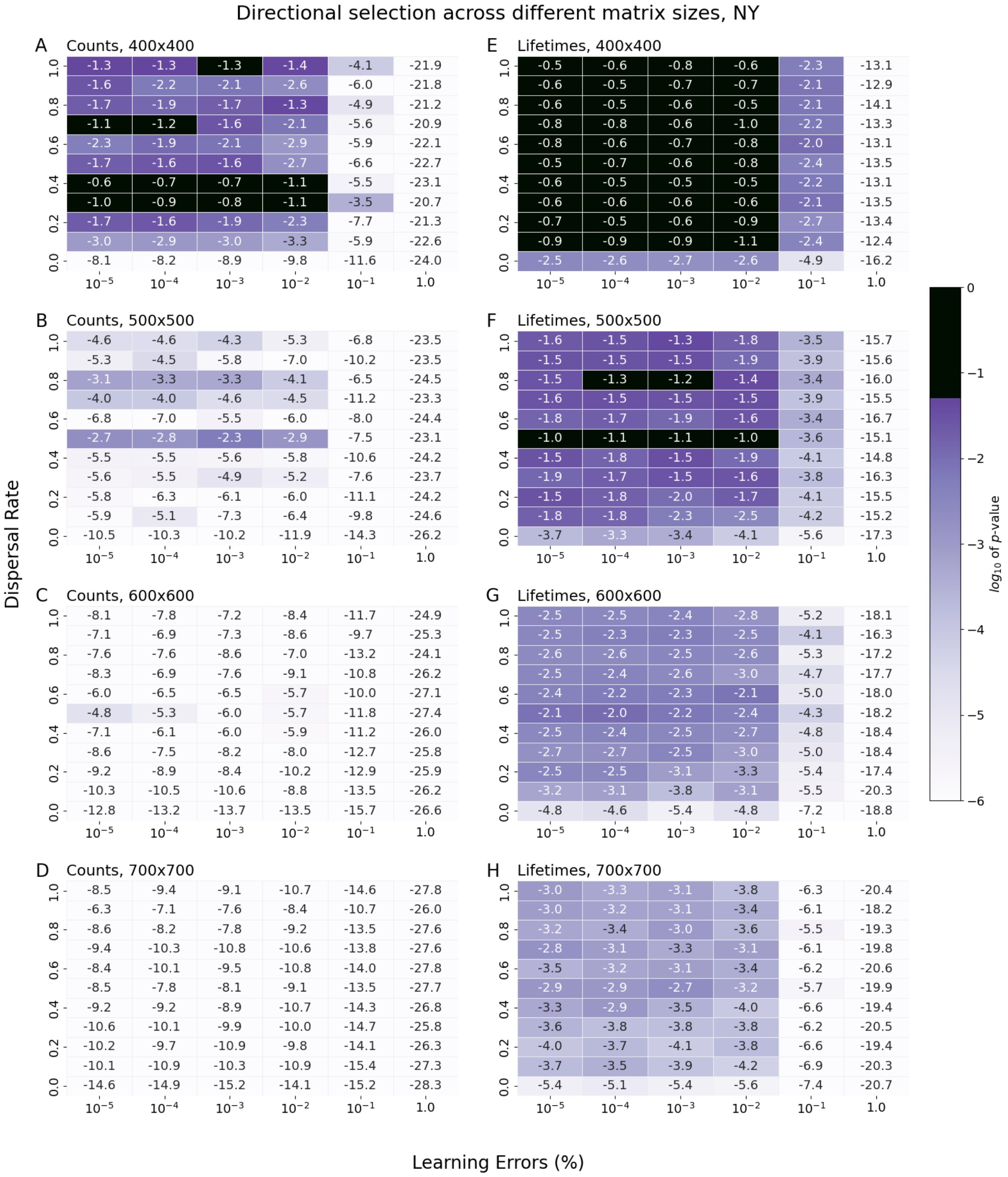


**Supplementary Figure S15. Comparisons of the effects of directional selection across different matrix sizes (400 by 400 to 700 by 700 in increments of 100), including counts (A-D) and frequency spectra (E-H) compared to empirical data in the NY region.** *P*-values for Fisher's exact tests in which the null hypothesis is that the binned syllable frequency spectra of empirical data and the simulated data are drawn from the same distribution. Models that are statistically indistinguishable ($P$>0.05) from the empirical data are shown in black. For neutral and conformity bias, see **Figs. S13-14**.

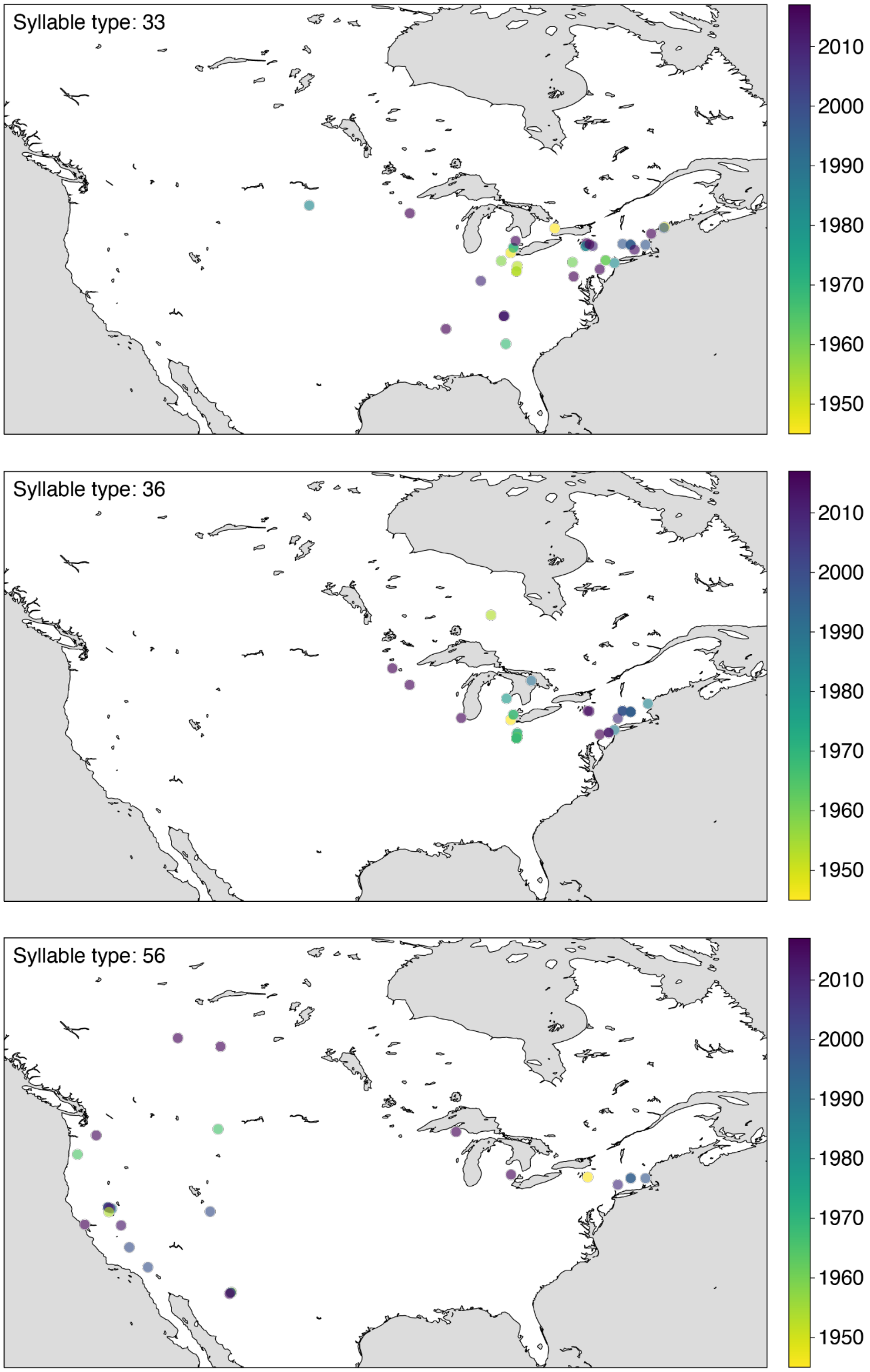


**Supplementary Figure S16. Maps showing the geographic and temporal distribution of three long-lived syllable types.** Syllable types 33 and 36 are found primarily in the eastern half of the United States, and syllable type 56 is found both in the east and west (see Searfoss, Liu, et al., 2020).
